# Single-cell sequencing of plasma cells from COVID-19 patients reveals highly expanded clonal lineages produce specific and neutralizing antibodies to SARS-CoV-2

**DOI:** 10.1101/2021.02.12.430940

**Authors:** Roy A. Ehling, Cédric R. Weber, Derek M. Mason, Simon Friedensohn, Bastian Wagner, Florian Bieberich, Edo Kapetanovic, Rodrigo Vazquez-Lombardi, Raphaël B. Di Roberto, Kai-Lin Hong, Camille Wagner, Daniel J. Sheward, Ben Murrell, Alexander Yermanos, Andreas P. Cuny, Miodrag Savic, Fabian Rudolf, Sai T. Reddy

**Affiliations:** Department of Biosystems Science and Engineering, ETH Zurich, Basel, Switzerland; deepCDR Biologics AG, Basel, Switzerland; Botnar Research Centre for Child Health, Basel, Switzerland; Department of Microbiology, Tumor and Cell Biology, Karolinska Institutet, Stockholm, Sweden; Institute of Microbiology and Immunology, Department of Biology, ETH Zurich, Zurich, Switzerland; Department of Pathology and Immunology, University of Geneva, Geneva, Switzerland; Swiss Institute of Bioinformatics, Mattenstr. 26, 4058 Basel, Switzerland; Department of Biomedical Engineering, University of Basel, Allschwil, Switzerland; Department of Surgery, Oral and Cranio-Maxillofacial Surgery, University Hospital Basel, Basel, Switzerland; Department of Health, Economics and Health Directorate, Canton Basel-Landschaft, Switzerland

## Abstract

Isolation and characterization of antibodies in COVID-19 patients has largely focused on memory B cells, however it is the antibody-secreting plasma cells that are directly responsible for the production of serum antibodies, which play a critical role in controlling and resolving SARS-CoV-2 infection. To date there is little known about the specificity of plasma cells in COVID-19 patients. This is largely because plasma cells lack surface antibody expression, which complicates their screening. Here, we describe a technology pipeline that integrates single-cell antibody repertoire sequencing and high-throughput mammalian display screening to interrogate the specificity of plasma cells from 16 convalescent COVID-19 patients. Single-cell sequencing allows us to profile antibody repertoire features in these patients and identify highly expanded clonal lineages. Mammalian display screening is employed to reveal that 37 antibodies (out of 132 candidates) derived from expanded plasma cell clonal lineages are specific for SARS-CoV-2 antigens, including antibodies that target the receptor binding domain (RBD) with high affinity and exhibit potent neutralization of SARS-CoV-2.

**One Sentence Summary:** Single-cell antibody repertoire sequencing and high-throughput screening identifies highly expanded plasma cells from convalescent COVID-19 patients that produce SARS-CoV-2-specific antibodies capable of potent neutralization.

## INTRODUCTION

Serum antibody responses against SARS-CoV-2 in convalescent COVID-19 patients play a critical role in controlling and resolving viral infection *(1)*. While SARS-CoV-2 consists of four major structural components, it is the multi-domain Spike (S) protein that represents the most relevant target for antibody-mediated immunity. The S protein is composed of the S1 subunit that includes the receptor-binding domain (RBD) and the S2 subunit which possesses target sites for the transmembrane serine protease (TMPRSS2). S1 is highly immunogenic and an important target during immunoglobulin (Ig) seroconversion and for viral neutralization, as the RBD directly interacts with the widely expressed human surface receptor angiotensin-converting enzyme 2 (ACE2), thereby initiating attachment and viral entry *(2, 3)*. Fusion of the viral particles with target cells requires a priming step which involves cellular proteases, such as TMPRSS2. Prevention of priming has also been shown to block SARS-CoV entry with a clinically-approved small-molecule inhibitor (Camostat) *(2, 3)*. Medical interventions such as infusion of SARS-CoV-2-specific antibodies through convalescent plasma from COVID-19 patients are being investigated and have received Emergency Use Authorization (EUA) by the United States FDA *(4, 5)* (NCT04383535 and CTRI/2020/04/024775). Furthermore, several therapeutic monoclonal antibodies originally derived from the B cells of COVID-19 patients and targeting the SARS-CoV-2 RBD have successfully progressed through clinical development, most notably bamlanivimab and etesevimab (Eli Lilly) which has also recently received EUA status *(3)*.

In addition to bamlanivimab *(6)*, several other antibodies originally discovered from COVID-19 patients are showing potential as therapeutics *(7–10)*, as highlighted by the rapid progression of several clinical trials (NCT04427501, NCT04452318, NCT04426695, NCT04519437). The isolation of human antibodies specific for SARS-CoV-2 have overwhelmingly focused on memory B cells (B_mem_), which due to their surface bound B cell receptor (BCR) can be labeled with soluble antigen (e.g., S1 or RBD) and sorted at high-throughput using flow cytometry, thus enabling selection and enrichment of antigen-binding B cells *(11)*. While integrated pipelines that incorporate single-cell sequencing are emerging to expedite antibody discovery *(12, 13)*, the isolation of antibodies from B_mem_ cells is still technically challenging due to a variety of factors such as the rarity of antigen-specific clones [e.g., 0.07% RBD-binding B_mem_ in COVID-19 patients *(14)*] and the requirement of single-cell sorting and cloning that can lead to dropout *(15)*. Another important consideration is that antibodies present in serum are not directly produced by B_mem_ but instead by antibody-secreting plasma cells (PCs).

Antigen-specific PCs can arise as early as within the first week of infection *(16)*, and thus can be profiled at an early time point of disease *(17)*. Recent studies have shown that many SARS-CoV-2-specific antibodies are very close to germline, often possessing stereotypical sequence patterns *(18, 19)*; with this in mind, it is possible that early or transitional PCs present in blood may encode specific and neutralizing antibodies to SARS-CoV-2. This transitional phenotype ranges from plasmablasts to short-lived PCs, prior to migration into the bone marrow and differentiation into long-lived PCs *(20)*. The frequency of transitional PCs in the peripheral blood mononuclear cell (PBMC) compartment of healthy donors is estimated at <5% of all B cells, but has been shown to increase to over 20% during COVID-19 infection *(21)*. This is a much more dynamic response than in B_mem_, where the fraction appears unchanged at 40-50% of total B cells during COVID-19 infection *(21)*. Previous research has shown that clonally expanded PCs following infection or immunization are often highly enriched for specificity to target antigens *(22–25)*; however it is yet to be determined if a similar phenomenon occurs in COVID-19 patients. A major challenge however is that in contrast to B_mem_, PCs cannot be pre-enriched for antigen binding due to low (IgA, IgM, IgE) or non-existent (IgG) surface expression of BCRs *(26)*.

In order to interrogate the specificity of PCs in convalescent COVID-19 patients, we describe here a technology pipeline that takes advantage of single-cell antibody repertoire sequencing, genome editing and high-throughput mammalian display screening (**Fig. 1**). We first perform single-cell antibody repertoire sequencing of PCs isolated from the blood of 16 convalescent COVID-19 patients. Single-cell antibody sequencing makes it possible to obtain information on the natural pairing of variable heavy (VH) and variable light (VL) chain sequences and to perform bioinformatic analysis to identify clonally expanded plasma cell lineages. Both the total repertoires as well as the expanded lineages show a broad germline gene usage and sequence diversity. We next select a total of 132 clonally expanded plasma cell lineages across all patients and design a CRISPR-Cas9-based genome editing strategy to rapidly integrate antibody genes into a mammalian display screening platform *(27–29)*. A selection approach based on fluorescence-activated cell sorting (FACS) combined with deep sequencing led to the discovery of 37 unique antibodies with specificity to SARS-CoV-2 antigens (S1, S2 and RBD). These antibodies are exclusively sourced from unique clonal lineages across 11 of the patients; notably there are three antibodies with specificity to the RBD that exhibit potent neutralization against SARS-CoV-2 in viral-pseudotype assays. Thus, our integrated workflow of single-cell sequencing and mammalian display screening is able to demonstrate that convalescent COVID-19 patients produce highly expanded PCs with specific and neutralizing antibodies for SARS-CoV-2.

**Fig. 1.**
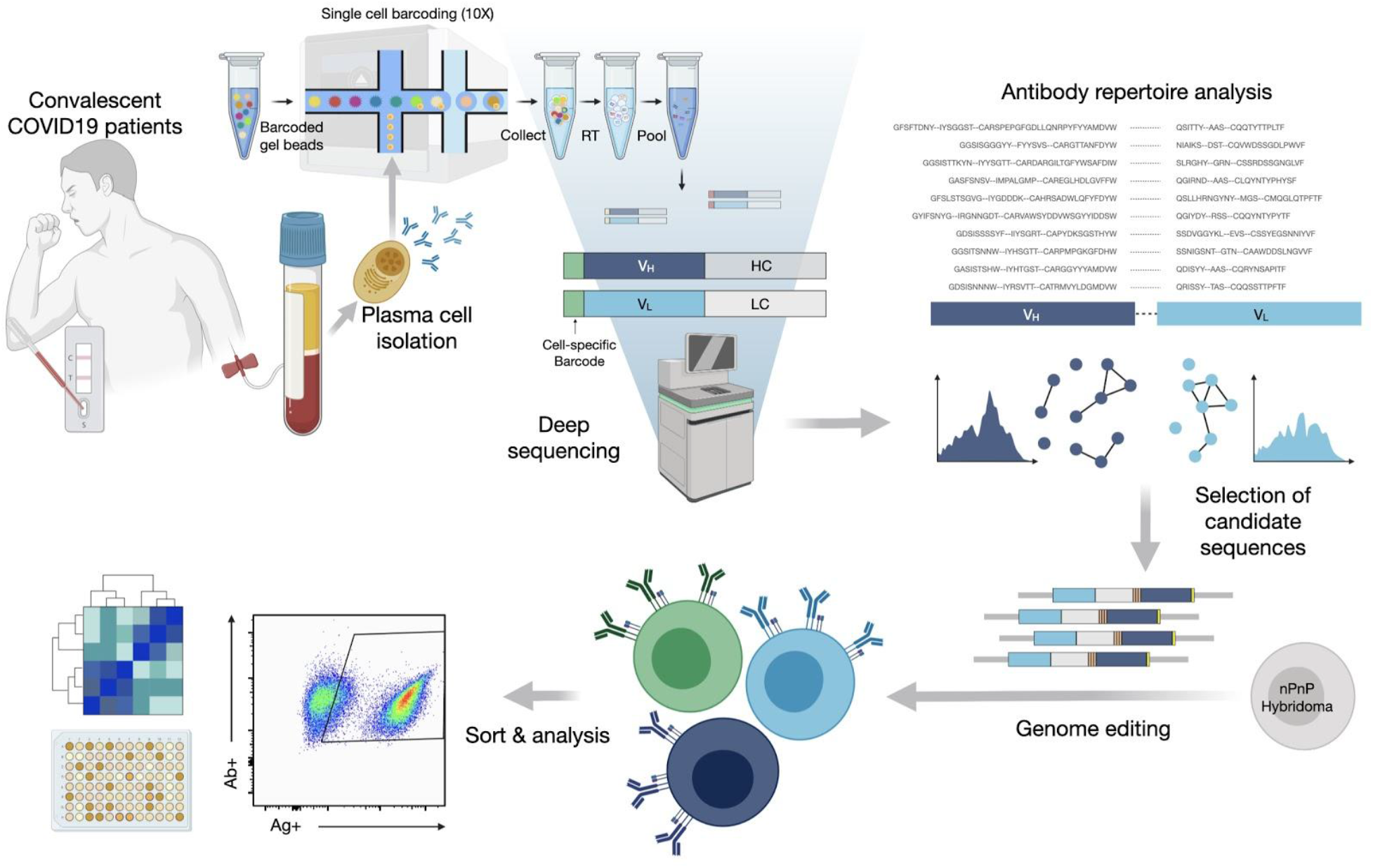
An integrated workflow for interrogating the antibody specificity of PCs from COVID-19 patients. Serum and peripheral blood mononuclear cells (PBMCs) are collected from convalescent COVID-19 patients (with confirmed PCR positive test). Serum is assayed with IgA and IgG ELISAs as well as POCTs. From a subset of 16 patients PCs are isolated from PBMCs by magnetic cell sorting and to then undergo gel encapsulation and barcoding for single-cell sequencing of their antibody heavy and light chain transcripts. Antibody repertoire analysis is performed to identify expanded plasma cell clonal lineages, which are then reformatted into single ORF full-length synthetic antibody genes including homology arms, to allow for single step cloning-free genome editing. The resulting mammalian display library then undergoes high-throughput screening for SARS-CoV-2 binding by flow cytometry and deep sequencing to recover the identity of the corresponding clonal lineages. Supernatant is used to determine cross-reactivity of antibodies within the library with coronavirus antigens.

## RESULTS

### A patient cohort of convalescent COVID-19 patients

Peripheral blood samples were isolated from a set of 16 convalescent COVID-19 patients, which was part of a larger cohort biobank (SERO-BL-COVID-19) *(30)*. Among the selected group of 16, all patients were confirmed to be SARS-CoV-2-positive by RT-PCR testing; point-of-care lateral flow tests (POCT) confirmed serum IgM and IgG antibody responses to SARS-CoV-2 S1 and nucleocapsid protein (NCP) antigens (**Fig. S1**). Clinical characteristics of the patients are summarized in **Fig. 2**; all patients displayed only mild symptoms (no hospitalization required) and with the exception of patients 2, 14 and 15, none of the patients required any assistance (**Fig. 2A**, left). Peripheral blood samples were collected within 9-24 days after onset of symptoms (average: 13 days) or 0-17 days after resolution of the disease (average ~5.1 days). A follow-up ELISA on patient serum (**Fig. 2B**, Euroimmun) failed to detect S1 specific IgA+IgG titers in two patients (F5921513, F5931620), and IgG titers in four patients (F5921407, F5921795, F5921393, F5921915), thus highlighting a discordance to POCTs. Sensitivity and specificity of the particular POCT used was >92% and 99%, respectively *(31)*.

**Fig. 2.**
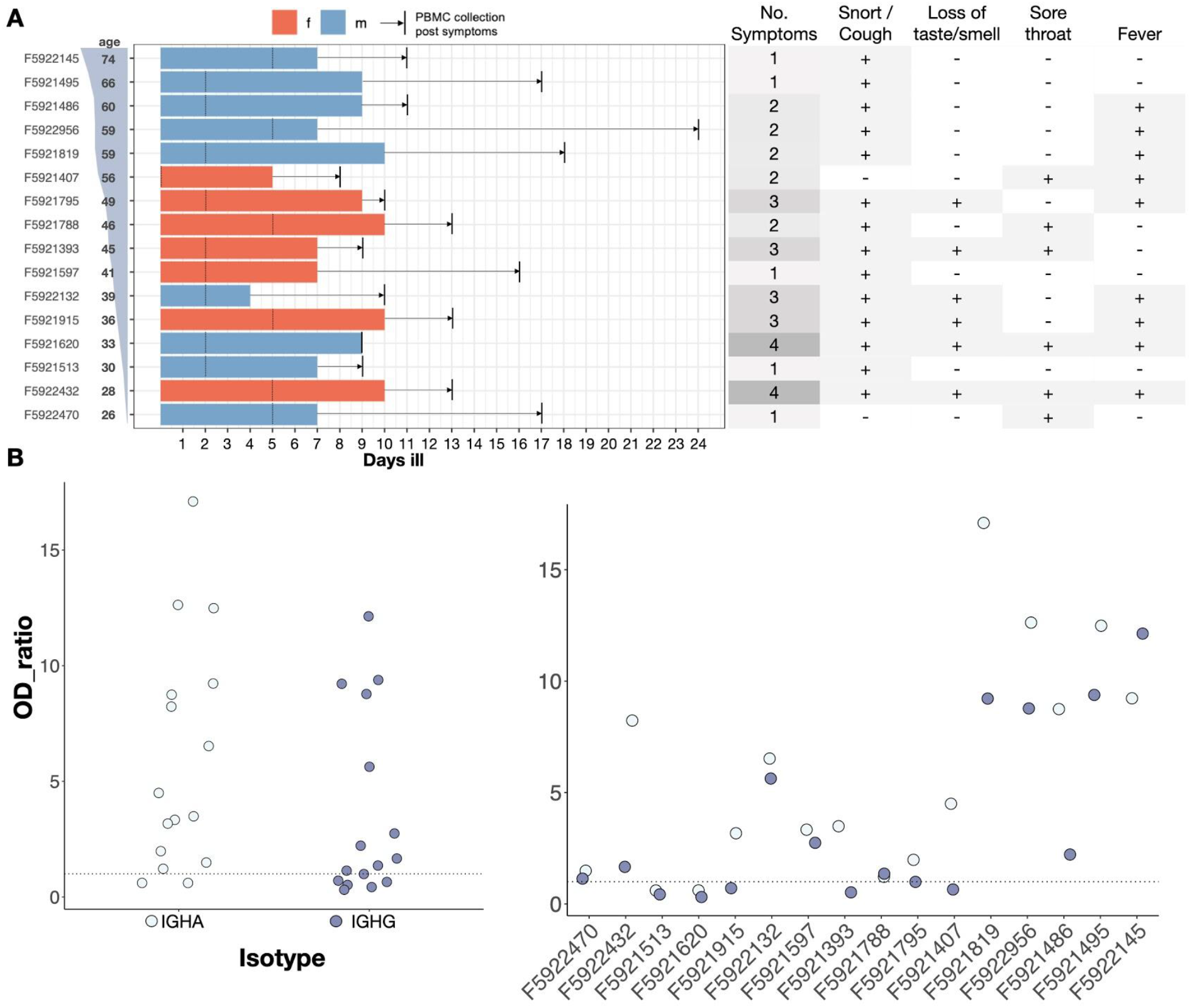
Demographics and symptoms of COVID-19 patients. A. Patients sorted by age. Bars, coloured by sex, show number of days ill total, before and after RT-PCR confirmed diagnosis (marked with a dotted line). Time between resolution of the disease and blood draw (vertical black line) is highlighted with an arrow. Summary of standardized symptoms per patient (right table). B. Euroimmun ELISA results over all patients (left) and per patient sorted by age (right) showing OD values over background for both IGHA (light blue) and IGHG (dark blue) titers. Dotted line represents classification cut-off.

### Single-cell antibody repertoire sequencing and analysis of PCs from convalescent COVID-19 patients

In order to interrogate the antibody repertoire of PCs in the cohort of convalescent COVID-19 patients, we performed single-cell sequencing. First, PCs were isolated from peripheral blood samples based on CD138 expression using magnetic beads. Next, single-cell sequencing libraries were prepared using the 10X Genomics Chromium system and the V(D)J protocol, which combines gel encapsulation and DNA-barcoding to tag antibody transcripts (mRNA) of both heavy and light chains originating from the same cell (see Materials and Methods for more details). Processing the raw reads using CellRanger (10x Genomics) resulted in approximately 8.3 million reads per patient plasma cell sample, with a mean depth of 176,000 reads per cell (**Table S1)**. Contigs were then assembled, filtered and re-aligned using Immcantation’s pRESTO/IgBlast pipeline *(32)*. Only cells with a productive heavy and light chain were considered for the final analysis. A plasma cell clone was defined by 100% amino acid identity of paired heavy and light chain complementarity determining region 3 (CDRH3-CDRL3) sequences. Clones with at least 80% similarity between their CDRH3 amino acid sequences and which possessed identical heavy chain germline genes were defined to be clonally related variants, forming a clonal lineage.

The number of cells where both heavy and light chain antibody sequences could be recovered varied across the 16 single-cell sequencing patient datasets, ranging from 30 to 1482 cells per sample (**Fig. S2A**). Analysis of the isotype usage across repertoires showed that the majority of clones were of IGHM (49.7±6.0%, mean±s.e.), although a substantial fraction of class-switched IGHG (29.6±4.4%) and IGHA (19.4±3.0%) was also observed (**Fig 3A**, **Fig S2B**). Next, we measured the percentage identity to germline V-gene, which served as a proxy for somatic hypermutation; this showed as expected that IGHG (94.4±0.1%) and IGHA (93.4±0.1%) had lower V-gene identity than IGHM (98.1±0.05%), however on a repertoire level this indicated that class-switched antibodies had only a minor degree of somatic hypermutation relative to repertoires from other viral infections (e.g., lymphocytic choriomeningitis virus) *(33)* (**Fig 3B**). We then examined the clonal expansion profiles per patient, identifying various degrees of clonal expansion, with several patients showing highly expanded IGHG and IGHA clones (**Fig 3C)**.

**Fig. 3.**
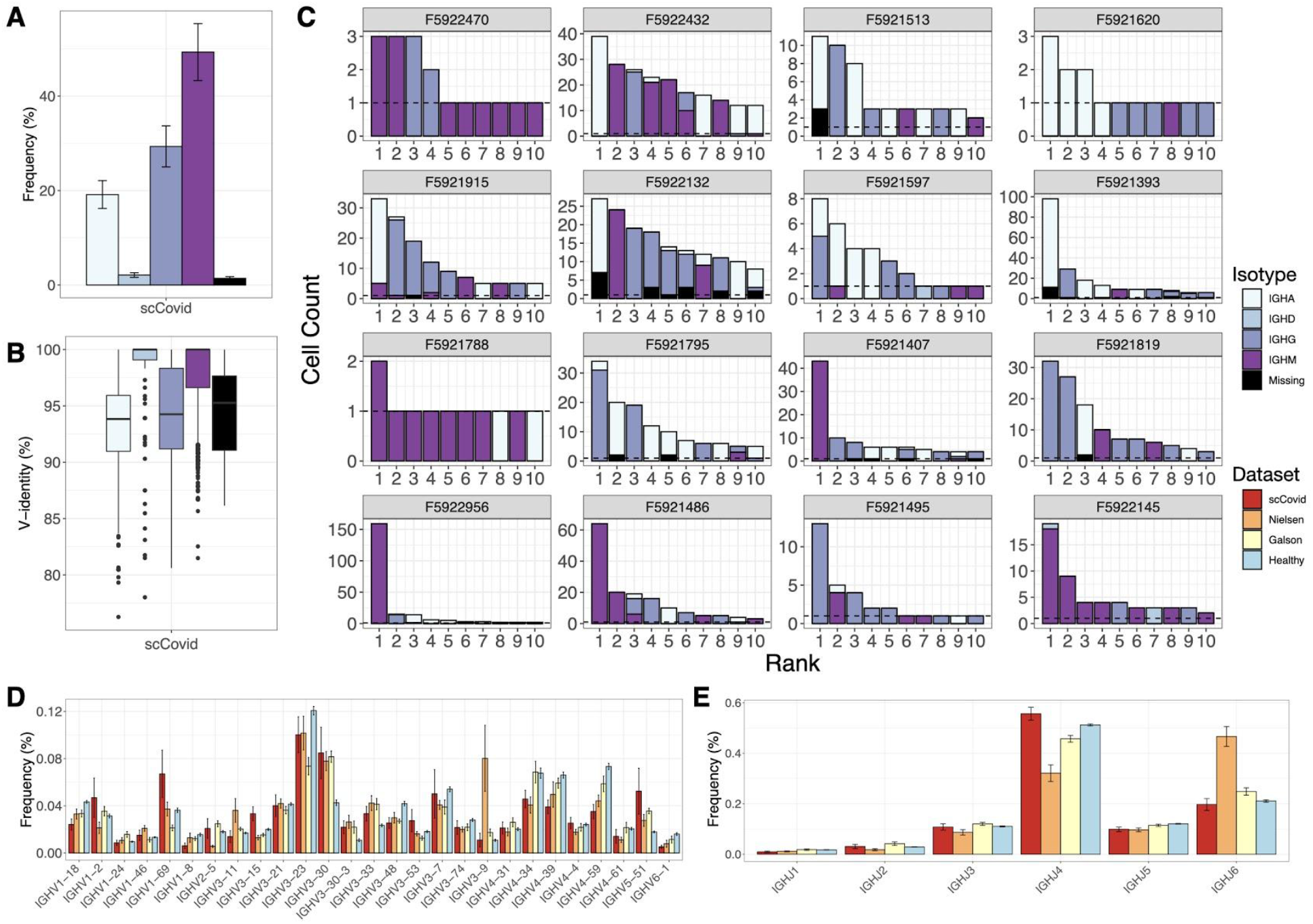
Single-cell antibody repertoire features and clonal expansion analysis. (A) Somatic hypermutation rates per isotype across all datasets. (B) Isotype usage in single-cell repertoires sequencing data of COVID-19 patients (error bars represent mean±se). (C) Clonal expansion profiles across all patients showing the top 10 most expanded clones, colored by isotype. (D) V and (E) J gene usage of heavy chains compared to previously reported repertoire sequencing datasets from COVID-19 patients and a healthy control.

Recently, papers from Nielsen et al. and Galson et al. describe antibody repertoire sequencing on B cells from COVID-19 patients *(34, 35)*. While these studies performed bulk sequencing (no linkage of natural pairing of V_H_ and V_L_) on peripheral B cells (mixture of B cell subsets) and did not interrogate antigen-specific binding, they do however offer a valuable reference of repertoire features such as germline gene usage during SARS-CoV-2 infection. We therefore compared the germline profiles from our single-cell repertoire data to the bulk repertoire data from these two studies and observed very similar trends in IGHV gene usage, including an increased usage of IGHV3-30, IGHV3-30-3, IGHV3-33 and IGHV5-51 relative to a healthy control repertoire data set *(36)* (**Fig 3D**). While an increased IGHJ6 usage was observed for the dataset from Nielsen et al., the overall usage distribution is similar for all datasets (**Fig 3E**). Analysis of the CDRH3 length distribution also showed no major difference across COVID-19 repertoire studies and the healthy control (**Fig S2A**).

Since our single-cell sequencing approach allowed us to recover heavy and light chain pairing information, we compared the pairing ratios of IGHV with IGKV or IGLV from our COVID-19 patients with that of pairing data from healthy individuals reported by DeKosky et al. *(37)*. The COVID-19 and healthy pairing ratios were generally very similar, notably IGKV3-20 and IGKV1-39 were both shown to pair with a wide range of IGHV, while others such as IGLV1-47 were only observed in combination with a limited number of IGHV (i.e. IGHV1-2) (**Fig. S2C**). Additional analysis of paired CDRL3 and CDRH3 length distributions did not reveal any trends or preferences in COVID-19 antibody repertoires.

### Selection of expanded plasma cell clonal lineages for mammalian display antibody screening

One of the major advantages of performing single-cell antibody repertoire sequencing was that it provided us with an opportunity to reconstruct full-length antibodies with natural V_H_ and V_L_ pairing, which allowed us to then interrogate the specificity of PCs against SARS-CoV-2 antigens. This approach requires the construction of synthetic antibody genes for recombinant expression, and thus one major limitation is that it is not cost effective to screen thousands of candidates. Although we implemented a streamlined design and workflow here, gene synthesis of thousands of antibody sequences became prohibitively expensive, thus emphasizing the importance of appropriate *in silico* analysis and selection. Despite the limitations in throughput, one of the critical questions that we could nevertheless answer is whether highly expanded plasma cell clonal lineages produced antibodies specific to SARS-CoV-2 antigens, as well as antibodies with neutralizing function. Therefore, we first started by selecting the IGHG clone with the highest abundance (cell count) in each of the 16 patients (**Fig. S3A**). We then selected an additional 20 antibody sequences associated with the most abundant clonal lineages from the four patients who had the highest single-cell sequencing depth (based on the number of cells with both V_H_ and V_L_ sequences recovered). This resulted in 36 unique antibody sequences and plasma cell clonal lineages, which were pooled together for subsequent antibody expression and screening experiments (Pool A). We selected and pooled an additional 96 antibody sequences associated with the most expanded clonal lineages across all 16 patients (Pool B). The resulting, highly diverse set of 132 candidate antibody sequences (**Table S2**) covered 104 unique heavy-light chain combinations (34 heavy and 30 light chains) (**Fig. S3B**) with CDRH3 and CDRL3 lengths ranging from 6 to 27 and 8 to 15 amino acids, respectively. The average germline identity to IGHV and IGKV/IGLV was 94.2% and 96.4%, respectively, including 26 variants that did not have any somatic hypermutations (100% germline identity). In order to efficiently screen the specificity of these antibody pools, we adapted our previously developed mammalian cell antibody surface display screening system: the plug-and-(dis)play (PnP) hybridoma platform, which facilitates the rapid generation of stable mammalian cell lines that display and secrete monoclonal antibodies *(27)*. PnP hybridomas take advantage of CRISPR-Cas9 and HDR to integrate a fluorescence reporter gene (mRuby) and subsequently synthetic antibody (sAb) genes into the endogenous IGHV genomic locus. Additionally, the stable integration of Cas9 into the genome and modification of HDR templates further improves integration efficiencies and enables the generation and screening of large mutagenesis libraries in specific regions such as the CDRH3 *(28)*, as well as entire variable gene libraries *(29)* and therapeutic antibody optimization by machine learning *(38)*. We started by adapting the PnP cell line by removing the fluorescent reporter mRuby from the V_H_ locus, thereby creating a negative PnP cell line (nPnP) (**Fig. S4A**). We hypothesized that this would improve HDR rates for templates with short homology arms, thus providing a cloning-free approach for antibody library generation and screening. Next, to design sAb templates for Cas9-induced HDR integration into nPnP cells, the following five gene modules were assembled: (i) the 5’ homology arm containing a partial V_H_1-82 promoter, followed by (ii) each V_L_ chain region (V-J) matched with its respective constant human light chain, either constant kappa (IGKC) or constant lambda (IGLC1), (iii) a flexible 57 amino acid linker, containing three tandem Strep-Tag(II) motifs providing both a connection to the heavy chain as well as a peptide tag for phenotypic detection of HDR integration and antibody expression *(39)*, (iv) the V_H_ chain region (V-D-J), followed by (v) a 3’ homology arm (consisting of the downstream intron). The full assembly of the modules resulted in a gene fragment of ~1.8 kb in size.

Commercial gene synthesis was employed to generate sAb HDR templates of Pool A and Pool B and the control CR3022, which is a previously identified cross-reactive antibody that binds SARS-CoV-1 *(40)* and SARS-CoV-2 and has and has high affinity (<0.1 - 6.3 nM) and specificity to a cryptic, non-neutralizing epitope on the S protein RBD *(41, 42)*. The sAb HDR templates and gRNA were co-transfected (electroporation by nucleofection) into nPnP cells and subsequent genotyping and phenotyping was performed. Following successful HDR mediated integration, mRNA from the sAb splices with the downstream IgH constant heavy chain (C_H_1) to assemble into a full-length antibody, thus only targeted integration results in antibody expression (**Fig. 4A,** left). Due to alternative splicing inherent to the endogenous IgH locus of PnP cells *(27)*, sAbs are both displayed and secreted into the supernatant **(Fig. 4A,** right). Cells displaying antibodies are thus derived from HDR-events, which can be selected by FACS when staining for anti-human IgG. The Strep-Tag is visualized on the surface by staining with Strep-Tactin (**Fig. 4B**). In order to screen a large number of antibody sequences at the same time, Pool A and Pool B were integrated into the genome and sorted. Enriched (HDR+) pools were then selected for binding to SARS-CoV-2 antigens, either to the S1 or S2 subunit, by staining the pools with both the tagged (mouse-Fc, mFc) antigens and a fluorescently labeled anti-mFc secondary, resulting in ~5-19% and ~2-7% binding populations, respectively (**Fig. 4C**).

**Fig. 4.**
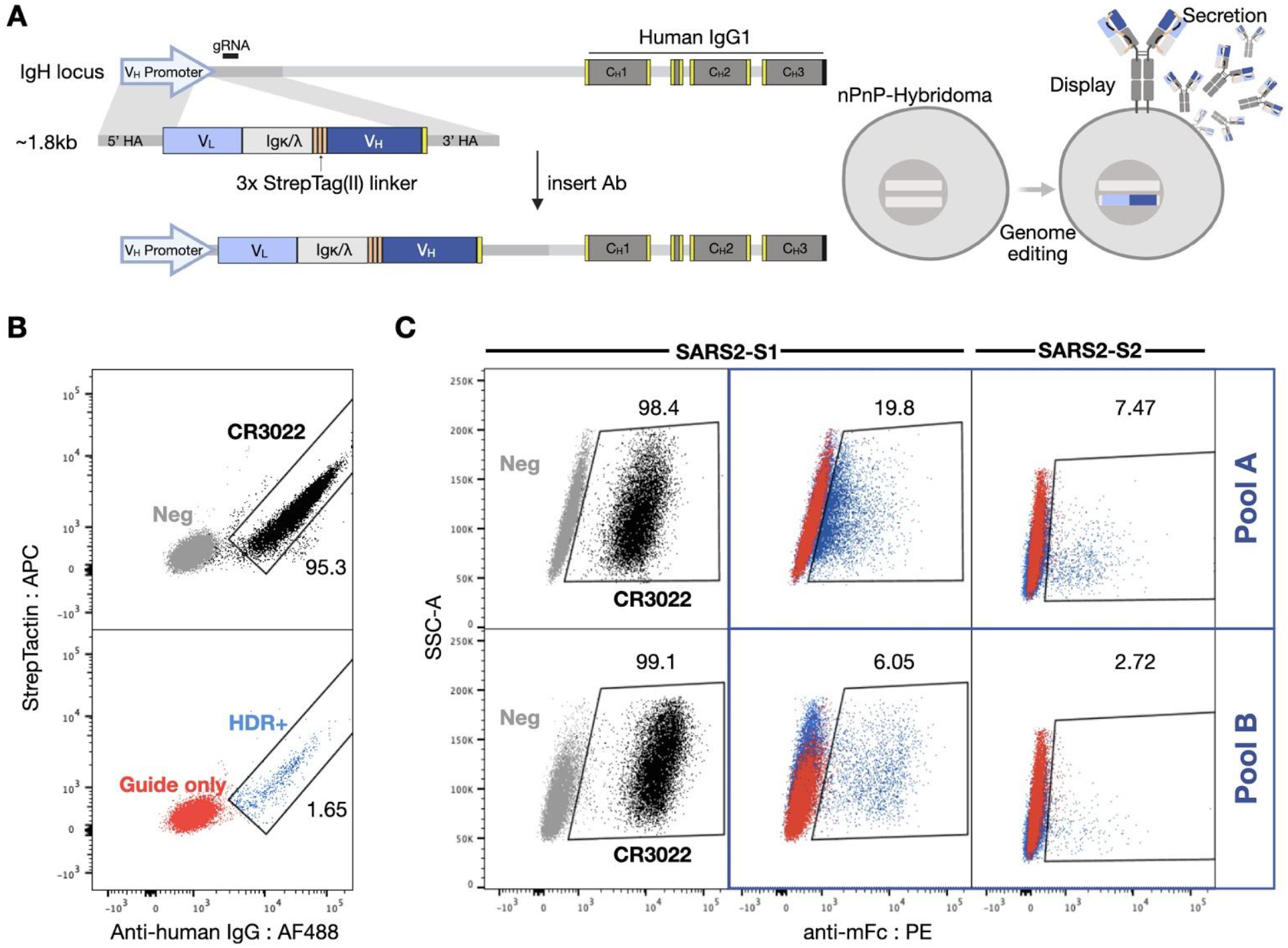
Mammalian display enables antibody screening of selected clonal lineages. (A) HDR templates are synthesized and transfected into PnP-hybridoma cell lines through CRISPR-Cas9 into the endogenous V_H_ locus. (B) After sorting by flow-cytometry for successful integration with Strep-Tactin and anti-human IgG, (C) enriched hybridoma pools (Pool A and B) are sorted for binding to SARS-CoV-2 S1 or S2.

### Validation of antibodies with specificity to SARS-CoV-2 antigens by deep sequencing and flow cytometry

In order to rapidly assess, screen and subsequently validate the specificity of antibodies towards SARS-CoV-2 antigens, we performed deep sequencing on Pool A and Pool B libraries following FACS. Genomic DNA was extracted from PnP cells of both pools after sorting for antibody expression (Ab+), as well as antigen binding (Ag+). Libraries were then prepared based on PCR amplification of the V_H_ gene (using a forward primer annealing at the 3x Strep-Tag and a reverse primer binding the start of the 3’ homology arm) (**Fig. S4B**). Samples were then subjected to deep sequencing (Illumina MiSeq v3 kit, 600 cycles, paired-end 2×300 bp), followed by alignment and post-processing using MiXCR *(43)*. Sequencing only the V_H_ region was still sufficient to determine the plasma cell clonal lineage, as all candidates that were chosen possessed unique combinations of CDRH1, H2 and H3. Hence, the sequence of the V_H_ region provides information to match it with the cognate V_L_ and thus recover the full candidate antibody sequence.

Out of the 132 total antibody candidate sequences present in both pools, 118 sequences (100% for Pool A, 85% for Pool B) were found in the deep sequencing data of the Ab+ population. Loss of some candidate sequences was likely a result of reformatting and the linker between variable chains, which may have prevented stable expression and assembly of antibodies in PnP cells. From the deep sequencing data, enrichment ratios were calculated by taking the log2 of the fraction of sequence counts in the Ag+ population and Ab+ population (**Fig. 5A, Table S3**). Highly enriched antibody sequences, as well as sequences differentially enriched for one of the S1 or S2 subunits were considered binders to the respective SARS-CoV-2 antigen and selected for downstream characterization. In addition, neutral or weakly depleted sequences were also chosen to help determine the enrichment cutoffs for binding vs. non-binding antibodies.

**Fig. 5.**
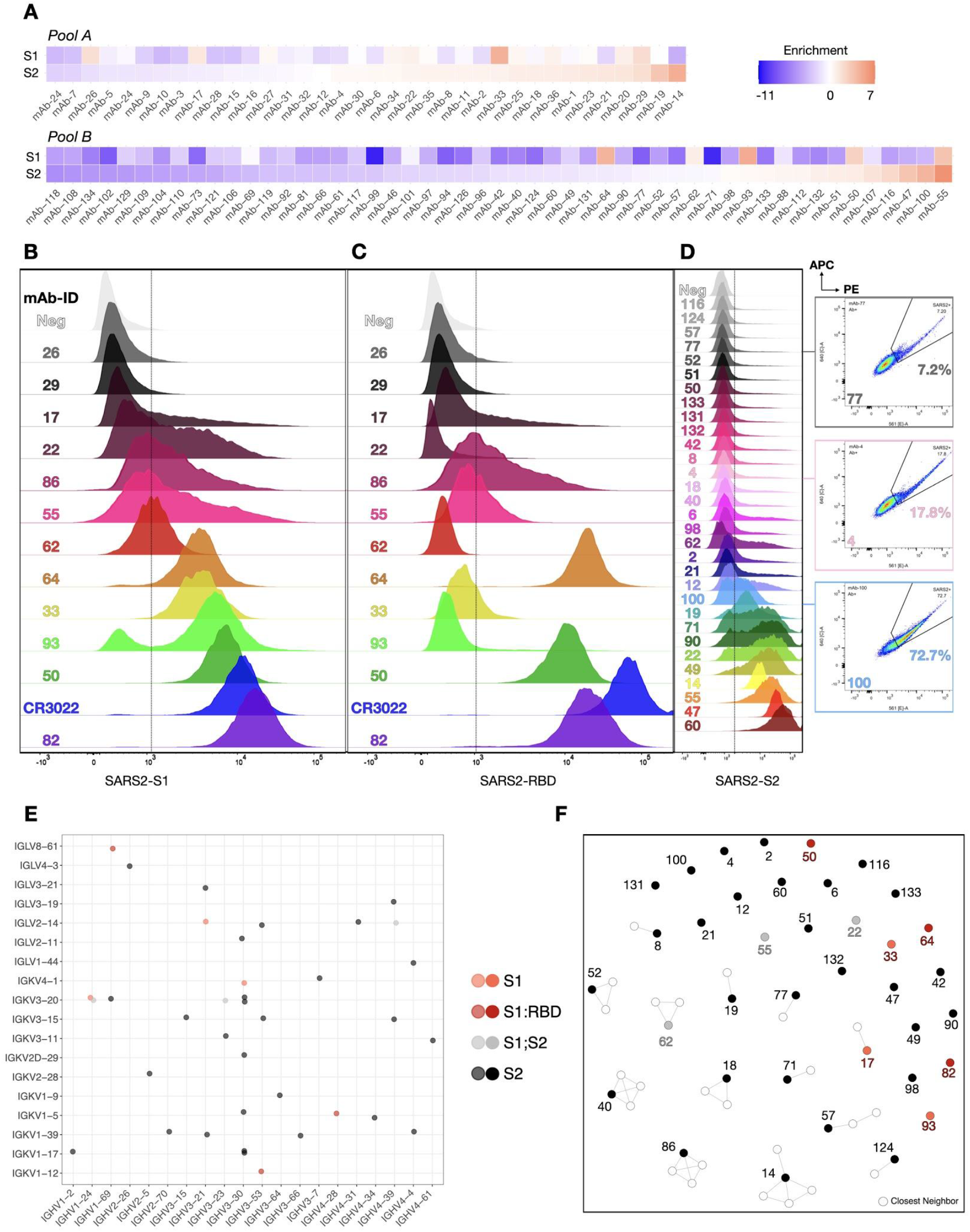
Validation of SARS-CoV-2 reactive antibodies by deep sequencing and flow cytometry. (A) Sorted cells are amplified using Strep-Tag-specific primers at 5’ and 3’ homology primers for Illumina Next-generation sequencing (see Fig. S4B). After MiXCR alignment, enrichment ratios are calculated to determine SARS-CoV-2 binders. Red tiles indicate highly enriched sequences, blue tiles correspond to strongly depleted sequences post FACS (B) RBD staining of S1 enriched candidates, (C) S1 staining of enriched candidates (D) S2 staining of enriched candidates, three example sequences (mAb-77, mAb-4 and mAb-100) shown on the side stained for both SARS-CoV-2 S2-PE and APC. (E) Dot-plot showing the V_H_/V_L_ gene usage across all verified reactive (S1, S1-RBD, S2) clonal lineages (F) Sequence similarity network of SARS-CoV-2 reactive sequences and their closest neighbors found in the patients repertoires (based on CDRH3 sequences, edges are drawn between sequences with Levenshtein distance ≤ 3).

Based on this enrichment data, a subset of 43 sequences was selected and Cas9 HDR was performed on nPnP cells to generate monoclonal antibody cell lines. Flow cytometry on the 43 monoclonal PnP cell lines was performed to verify if antibodies were specific to SARS-CoV-2 S1, RBD or S2 antigen **(Fig. 5B-D)**. Out of 43 tested candidates, 37 displayed strong or sufficient Stokes shifts in the fluorescence channel (561 nm) to verify their specificity. For some candidates, strongly trailing peaks indicated an incomplete staining possibly due to differential antibody expression and stability. Nevertheless, these antibodies can still be considered specific to SARS-CoV-2 antigen due to a fluorescence signal in both 561 and 670 nm channels (**Fig. S5A**). Additionally, an unrelated mFc-tagged antigen (same isotype, IgG1) (here: anti-hCD69-mFc) was used to verify that the observed signal was not a result of mFc interactions (selected samples remained negative for the unrelated mFc-tagged antigen, while positive for S1 and S2 antigens) (**Fig. S5B**). Candidate antibodies with specificity to S1 (**Fig. 5B**) were also labeled with the RBD antigen, revealing that at least three clones appeared to bind to the RBD **(Fig. 5C)** and could potentially be neutralizing antibodies. Flow cytometry confirmed that most of the highly enriched clones, but surprisingly also some differentially depleted clones, showed specificity to SARS-CoV-2 antigens, with varying degrees of intensity [mean fluorescence intensity (MFI) = 383-17,165 for S1 (background = 388), MFI = 376-65,396 for RBD (background = 354) and MFI = 923-50,408 for S2 (background = 826)] **(Fig. 5D)**. Both S1- and S2-specific clones displayed a breadth of signal, most likely derived from the dynamic range of antibody surface expression on PnP cells and a large range of affinities.

In order to investigate any potential cross-reactivity to other common human coronaviruses (CoV), supernatant from Pool A and Pool B was collected and ELISAs were performed with antigens derived from four other CoV: 229E, HKU1, OC43 and NL63, in addition to SARS-CoV-2 antigens S1, S2 and nucleocapsid protein (NCP) (**Fig. S6**). As a reference, the CR3022 supernatant showed strong binding to S1 and S2 antigens, while showing no cross-reactivity with any other CoV antigen. The antibodies from Pool A showed more binding to all CoV antigens, while the Pool B antibodies appeared to be more focused on S1 and NCP antigens, which was correlated with what was observed in FACS experiments.

### COVID-19 patient and SARS-CoV-2-specific antibody sequence analysis

Based on these measures, 37 of the 132 (28%) antibody sequences selected from the highly expanded plasma cell clonal lineages could be confirmed to be specific for SARS-CoV-2 S1, RBD or S2 antigen (**Table S4**). By examining these SARS-CoV-2-specific sequences and which COVID-19 patients they originated from, we observed patient-related trends (**Table S5**). For instance, 44% (4/9) of antibody sequences selected from patient F5921407 were SARS-CoV-2 antigen-specific, while for another patient only 10% (1/10) of tested antibody sequences demonstrated antigen binding. As the number of selected sequences varied per patient, the mean hit-rate per patient was slightly higher at 35%. From our cohort, serum from two of the patients (F5921513, F5921620) showed a complete lack of S protein reactive antibodies by ELISA (**Fig. 2B**), which is in stark contrast with previous studies showing all COVID-19 patients develop some S protein-specific antibody response *(44, 45)*. To further corroborate this, none of the four antibody sequences (mAb-1, mAb-9, mAb-104 and mAb-135) from highly expanded PCs of these two patients were shown to bind to S1 or S2 antigens. The 37 observed SARS-CoV-2-specific antibody sequences were distinct from each other and did not strongly correlate with trends observed in the overall repertoire. For example, there were 34 unique V_H_-V_L_ germline combinations and a broad distribution of V- and J-gene usage (**Fig. 5E, Fig. S7A)**. The most frequently observed VH-gene (IGHV3-30, seen in 8 binders) was also present at an increased frequency in both the overall repertoires as well as the pool of 132 selected sequences. The CDRH3 lengths ranged from 6 to 26 (CDRL3: 8 and 15) **(Fig. S7B)** and the set of binders exhibited a broad range of somatic hypermutation with V-gene identities between 85%–100% **(Fig. S7C)**. Notably, 11 of the antibodies possessed no somatic hypermutations, with a germline identity of 100% in both their V_H_ and V_L_ sequences. Comparing our results to previously published SARS-CoV-2-specific antibody sequences, we found that two clones, mAb-64 and mAb-93, were present in a similar form in other available datasets *(35, 46)*. Specifically, mAb-64 has six clonal neighbors with ~85% similarity in the CoV-AbDab (a database compiling known coronavirus-binding antibody sequences) *(46)*, while the CDRH3 sequence of mAb-93 was also found in the previously published COVID-19 repertoire dataset from Galson et al. *(35)*. Notably, these SARS-CoV-2-specific antibody sequences only have a limited number of close neighbors (Levenshtein distance ≤ 3) in our COVID-19 patient repertoires (**Fig. 5F**). The sparsity of the similarity network though is likely in large part due to the limited size of the single-cell sequencing datasets.

### Clonally expanded PCs produce high affinity antibodies to the RBD that are capable of neutralizing SARS-CoV-2

Of special interest are the PCs and their corresponding antibodies with specificity to the RBD, as binding to the RBD is often associated with neutralization of SARS-CoV-2. The three antibodies in this group used different combinations of V_H_ and V_L_ germline genes, namely IGHV4-28-IGKV1-5 (mAb-50), IGHV3-53-IGKV1-12 (mAb-64), IGHV1-69-IGLV8-61 (mAb-82). Interestingly, IGHV3-53 and IGHV1-69 have previously been reported in association with several other neutralizing antibodies to SARS-CoV-2 *(9, 18, 47–49)*. Previous work has established that B_mem_ can produce antibodies that are germline or have very few somatic mutations and be specific for SARS-CoV-2 antigens, including the RBD *(19)*. Thus, similar to B_mem_, we found this similar trend in PCs, as two of the three antibodies (mAb-64, mAb-82) were completely germline sequences in both V_H_ and V_L_ chains, while the third (mAb-50) had a V-gene identity of 94%, which was the same as the average for the whole repertoire. Next, we measured the binding affinities of the RBD-specific antibodies by using a biotinylated prefusion stabilized SARS-CoV-2 S1 antigen and biolayer interferometry (BLI). All three antibodies exhibited similar binding kinetics, with apparent equilibrium dissociation constants (*K*_*D*_) values of 1.61 nM (mAb-50), 1.96 nM (mAb-64) and 1.78 nM (mAb-82) (R^2^ = 0.99). While mAb-50 exhibited the fastest on-rate (6.80 M^−1^s^−1^), approximately 3.5 times faster than either mAb-82 (2.05 M^−1^s^−1^) or mAb-64 (1.89 M^−1^s^−1^), it possesses an off-rate that is faster by about the same factor (**Fig. 6A-C**). Antibodies derived from B_mem_ have thus far been reported to routinely display affinities in the nanomolar (*K*_*D*_ ≈ 1 × 10^−9^) to subnanomolar (*K*_*D*_ < 1 × 10^−9^), and even in some cases the low picomolar range (*K*_*D*_ ≈ 1 × 10^−12^) *(7, 13, 14, 19, 26, 50)*. While the RBD constitutes only a small subsection of the S1 antigen, it contains at least four distinct epitopes *(51)* for antibody binding, which can potentially impact the capacity for neutralization in different ways. Targeting multiple different (non-overlapping) epitopes has been shown to potentiate neutralization and prevent viral escape in SARS-CoV-1 and, recently, SARS-CoV-2 *(40, 52)*. Therefore, we performed epitope mapping of our plasma cell-derived antibodies by using an in-tandem competition assay: SARS-CoV-2 S1 was immobilized to Streptavidin biosensors, followed by incubation with an antibody 1 and a secondary, competing antibody 2 at a lower concentration. As controls we used the CR3022 antibody or the extracellular domain of the ACE-2 receptor (**Fig. 6D**). As expected, competition and self-blocking occurred between pairs of the same antibody. None of the plasma cell antibodies competed with CR3022, but all antibodies were able to block the binding of a secondary ACE2. For CR3022, the binding intensity of the secondary antibody was found to be approximately half of the competition free response, indicating a weak interaction or overlap. Given that CR3022 binds a cryptic epitope that requires a conformational change of the RBD *(41)*, binding of another antibody may reduce the stability of such a change. The germline-like antibodies mAb-64 and mAb-82 displayed similar behaviour, and were unable to bind in the presence of one another, thus indicating an overlapping epitope. The epitope targeted by mAb-50 did not overlap with either mAb-64 or mAb-82 (**Fig. 6E**). Finally, to determine if RBD-specific antibodies were indeed able to neutralize SARS-CoV-2, we performed neutralization assays using HEK293T-ACE2 cells and SARS-CoV-2 pseudotyped lentivirus *(53)*. An alpaca-derived single domain antibody (ty1) was used as a positive control for neutralization *(53)* (**Fig. 6F**). IC_50_ values were determined to be 145.0 ng/mL (mAb-50), 104.7 ng/mL (mAb-64), 120.2 ng/mL (mAb-82), while neutralization saturates below 100% at 93.4%, 95.9% and 96.1%, respectively. These results verify that transitional PCs in blood are indeed antigen specific and demonstrate that single cell sequencing on PCs can be utilized to select and identify high affinity and potently neutralizing antibodies against SARS-CoV-2.

**Fig. 6.**
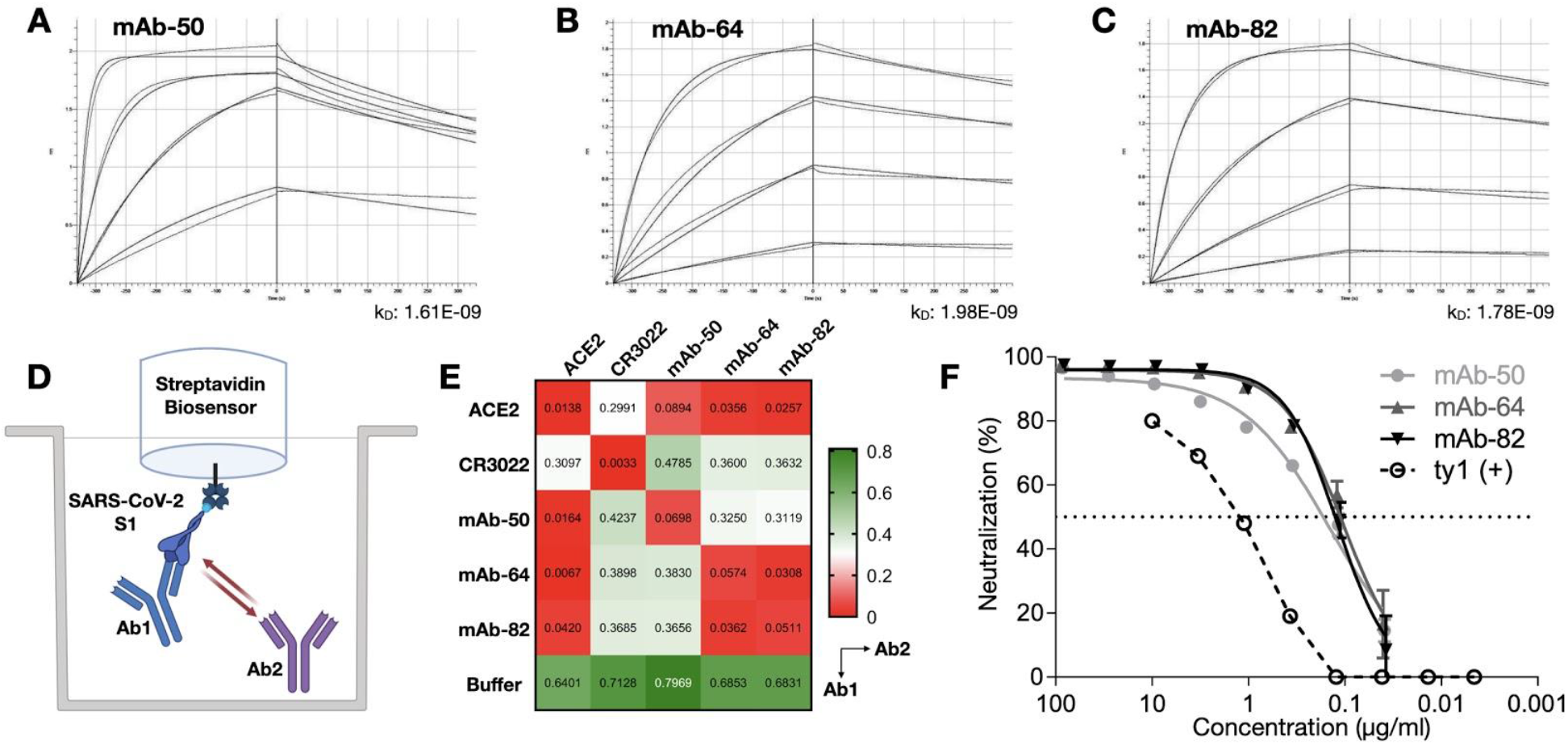
Characterization of RBD-specific antibodies for affinity, epitope binning and neutralization. (A-C) Antibodies were purified from supernatant and assayed at 5 different concentrations ranging from approximately 2 to 500 nM [shown here: 125 nM (green), 37.5 nM (yellow), 7.8 nM (purple), 1.9 nM (teal)]. Software calculated fits are shown in red. Binding kinetics (apparent affinity constant kD) to SARS-CoV-2 S1 for FACS-confirmed RBD binders, determined by bio-layer interferometry. (D) Binders were subsequently assayed to perform epitope binning by immobilizing SARS-CoV-2 S1 on the sensors, followed by a first antibody at 166 nM, followed in tandem with a secondary antibody at 78 nM. (E) Additional binding by the secondary molecule indicates an unoccupied epitope (non-competitor, green), while no binding indicates epitope blocking (competitor, red). Self-blocking confirmation can be found on the diagonal. (F) Inhibition of infection of HEK293T-ACE2 cells with SARS-CoV-2 pseudotyped lentivirus.

## DISCUSSION

Here, we elucidated the antibody response of PCs in convalescent COVID-19 patients by performing single-cell antibody repertoire sequencing combined with mammalian display antibody screening. Single-cell sequencing revealed a feature landscape that was highly consistent with previously reported COVID-19 patient repertoires with respect to germline gene usage and CDR3 length distribution *(34, 35)*. We also observed a comparatively low rate of somatic hypermutation for Ig class-switched antibodies on a repertoire level, a finding in line with previous reports on antibodies associated with SARS-CoV-2 infection *(7, 34)*. Comparing sequences of our COVID-19 patient repertoires with sequences from the CoV-AbDab, we were able to identify 45 antibodies with an amino acid sequence similarity above 80% in CDRH3 **(Table S6)**. In addition, we also found that mAb-64 shared high sequence similarity (up to 92%) to recently described stereotypical sequences of RBD-binding antibodies found to be commonly present in COVID-19 patients *(18)*. The recovered heavy and light chain pairing information was in line with previously reported data from healthy donors, with an increased frequency of common heavy and light pairs. IgG isotype-resolved lineage expansion analysis enabled the identification of 132 highly expanded plasma cell clonal lineages, representing a diverse sequence space, to be tested for specificity to SARS-CoV-2 antigens.

In order to enable the rapid screening and discovery of SARS-CoV-2 reactive candidates within highly expanded plasma cell lineages, we adapted a mammalian display platform based on the previously reported PnP-hybridoma system *(27)*. In a cloning-free, CRISPR-Cas9 HDR-mediated process, we created two antibody libraries (Pool A and Pool B) and screened for binding to SARS-CoV-2 antigens by flow cytometry. Through the use of deep sequencing and enrichment analysis, we could effectively determine the identity of antibodies binding to SARS-CoV-2. A subset of these antibody candidates were then integrated individually into the PnP system to generate monoclonal cell lines in order to verify and further characterize specificity to SARS-CoV-2 antigens. Thereby, we could show that in most of the patients in our cohort (11/16) there were highly expanded PCs producing antibodies with specificity to SARS-CoV-2. While we were unable to discover antibodies from all patients with specificity to S1 or S2, this could be explained by the fact that some patients had low or no detectable serum antibody titers against S protein (**Fig. 2B**) or in some patients there was low single-cell sequencing depth that may have compromised clonal expansion analysis (**Fig. 3C)**. In both Pool A and Pool B, antibodies showed cross-reactivity to other human CoV antigens, including HCoV-229E, OC43, HKU1 and NL63, but did not show any specificity to MERS-CoV-Spike or SARS-CoV-1 Spike, which is in contrast to the cross-specific antibodies discovered by others *(8, 9)*, in which a large fraction of candidates potently bound both SARS-CoV-1 and SARS-CoV-Importantly, Pinto and colleagues successfully screened B cells from recovered SARS-CoV-1 patients for SARS-CoV-2 binding, proving that certain epitopes are preserved; Wec et al. followed a slightly different approach, where a convalescent COVID-19 patient was chosen that had also survived a SARS-CoV-1 infection in the 2003 SARS epidemic. While SARS-CoV-1 patients thus seem to contain SARS-CoV-2 reactive antibodies, other discovery efforts from COVID-19 patients (that did not include SARS-CoV-1 experienced donors) have been unable to discover antibodies with cross-reactivity *(54)*. However, a recent study by Walker and colleagues that applied protein mutagenesis and directed evolution suggests it is possible to engineer SARS-CoV-2-specific antibodies to have cross-reactivity and neutralization across a large panel of human CoV variants *(55)*.

Similar to the previous study by Ju and colleagues, there were no generalizable repertoire or sequence features within the discovered SARS-CoV-2-specific antibodies, further supporting the diversity of antibody repertoires that develop in COVID-19 patients. Despite this heterogeneity, we did find sequences that appear to be similar to those already reported *(14, 35)*, hinting at the importance of a germline-driven response in reaction to an infection with SARS-CoV-2 *(19, 47)*. The overall number of antibodies that were identified to be specific for SARS-CoV-2 in this study is likely an underestimation, as not all of the pooled candidates enriched by flow cytometry and deep sequencing were tested separately. Based on the 81% success rate (35/43) of individually expressed antibodies, we believe that a number of yet untested candidates are likely to be positive for SARS-CoV-2 binding. Additionally, antibodies were only screened for binding to the S protein subunits, while there are possibly a number of antibodies that also bind to the NCP antigen based on ELISA reactivity of the Pool A supernatants (**Fig. S6**). Candidate sequences that were missing from the library based on deep sequencing data potentially had stability and expression issues in the PnP system. While the linker-based design of the HDR construct improved and facilitated the generation of stable cell lines, reformatting does come with its limitations similar to known issues of converting antibody fragments (e.g., single-chain variable fragments and fragment antigen binding) to full-length IgG *(56, 57)*, as well as affinity, stability and expression issues. Evaluating the RBD binders in the non-linker format increased the affinity and neutralizing potential, emphasizing the negative influence of a linker on antigen binding. This limitation may have caused a substantial loss in SARS-CoV-2 reactive sequences that do not tolerate the presence of the utilized 57-amino acid linker.

Recently, Cao and colleagues have demonstrated how clonal expansion of B_mem_ can be used as a proxy for antigen specificity *(13)*. However, despite finding two RBD-specific antibodies among the 132 candidates that were expressed, only one of them appeared to be (weakly) neutralizing, leading to the conclusion that antigen pre-selection was required. In this study, we found three potent neutralizing antibodies targeting the RBD from 132 selected clonal lineages, and while this is only a slight improvement in the success rate compared to Cao et al., it did demonstrate the potential to discover antibodies from PCs by single-cell sequencing. Given the unique physiology of PCs, this is particularly relevant at early stages of an infection as well as in the first weeks and months of a pandemic, when expediency is required and antigen-based selection may not be possible due delayed availability of antigen or reduced B_mem_ formation. It is important to note that serum antibodies represent an exceptional source for clinical antibody candidates *(58)* to which plasmablasts, short- and long-lived PCs are the sole contributors *(17)*. In the future, different strategies of selecting plasma cell clonal lineages could allow for improvement in successful identification of antigen-specific antibodies from single-cell sequencing data. In addition to clonal expansion, other factors could be incorporated into the selection criteria such as Ig isotype, somatic hypermutation and convergence in sequence space *(59)*.

## MATERIALS and METHODS

### Study design

Here, we set out to profile and characterize SARS-CoV-2 specificity in PCs obtained from convalescent COVID-19 patients with positive Point-of-Care lateral flow test for N/S protein (Qingdao Hightop Biotech, #H100). Non-hospitalized patients older than 18 with a normal immune system and positive RT-PCR test were derived from the Swiss project “COVID-19 in Baselland Investigation and Validation of Serological Diagnostic Assays and Epidemiological Study of SARS-CoV-2 specific Antibody Responses (SERO-BL-COVID-19)”, as reported earlier *(30)*.

The study used single-cell antibody repertoire sequencing of patient PCs and the analysis of the resulting repertoire. Expanded lineages were selected for further investigation by synthesis and CRISPR-Cas9 mediated stable integration into the PnP hybridoma-derived mammalian display platform. Flow cytometry aided in selecting antigen-specific cells performed in high-throughput, followed by deep sequencing to identify the antigen-specific sequence by enrichment. SARS-CoV-2 S1-, RBD-and S2-specific antibodies were evaluated individually by flow cytometry with mouse-Fc tagged antigens, followed by a secondary antibody tagged with Phycoerythrin (PE) or Allophycocyanin (APC). RBD-specific antibodies were directly Strep-Tag(II)-purified from supernatant of monoclonal cell lines and assayed on Octet as well as viral neutralization assays.

### Ethical statement

Ethics permission was granted by the ethics board “Ethikkommission Nordwest- und. Zentralschweiz (EKNZ)”. Every participant has signed a written informed consent as described previously *(30).*

### Patient samples

In April 2020, a study on the detection accuracy of POCTs was launched by the Department of Health of Basel-Land and Basel-Stadt, testing COVID-19 patients with mild symptoms and positive RT-PCR test with POCTs from a number of manufacturers. Over 300 patients were characterized and recruited for PBMC and serum collection.

Participants consisted of patients from the SERO-BL-COVID-19 study sponsored by the Department of Health, Canton Basel-Land, Switzerland. SARS-CoV-2 infection was confirmed by RT-PCR of nasopharyngeal swab samples. Importantly, participants had mild to moderate symptoms without hospitalization. Patients included in the subset were selected based on POCTs assessing the presence of IgG and IgM SARS-CoV-2-specific antibodies (Qingdao Hightop Biotech, #H100), performed at the time of blood collection, after resolution of symptoms as seen from **Fig. 1**. PMBCs were isolated as described previously *(30)*. In brief, PBMCs were isolated from EDTA blood using density gradient separation in Ficoll Paque Plus reagent (GE Healthcare, #17-1440-02). After separation, plasma was collected for ELISA detection of IgG and IgA SARS-CoV-2-S1 specific antibodies (Euroimmun Medizinische Labordiagnostika, #EI2668-9601G, #EI2606-9601A). PMBCs were then resuspended in freezing medium (RPMI 1640, 10%(v/v) FBS, 10%(v/v) dimethyl sulfoxide) and cryopreserved in liquid nitrogen.

### Immunomagnetic isolation of PCs

PBMC samples were thawed, washed in complete media (RPMI 1640, 10%(v/v) FBS), followed by centrifugation. The cell pellets were resuspended in 0.5 mL complete media, counted and treated with 10 U mL^−1^ DNase I (STEMCELL Technologies, #07900) for 15 min at RT to prevent cell clumping. Cells were then washed again and pelleted by centrifugation. After resuspension in 0.5 mL flow cytometry buffer (PBS, 2%(v/v) FBS, 2 mM EDTA), the cells were filtered through a 40 μm cell strainer prior to immunomagnetic isolation, using an EasySep Human CD138 Pos. Selection kit II (STEMCELL Technologies, #17877) according to the manufacturer’s instructions. Whenever possible, cells were adjusted to a concentration of 1×10^6^ live cells/mL in PBS, 0.04%(v/v) BSA before proceeding with droplet generation.

### Single-cell antibody repertoire library preparation

PCs were encapsulated with DNA-barcoded gel beads using a 10x Chromium controller (PN-110203). Samples were counted using a BioRad TC-10 automated cell counter and normalized to 1.7 ×10^4^ cells in reverse transcription (RT) mix to ensure the recovery of 1 ×10^4^ cells post-loading. After recovery, target enrichment RT and library preparation was performed according to the manufacturer’s instructions (CG000086 manual, RevM, 10x Genomics) using the following kits: Chromium Single Cell 5’ Library & Gel Bead Kit (PN-1000006), Chromium Single Cell 5’ Library Construction Kit (PN-1000020), Chromium Single Cell V(D)J Enrichment Kit, Human B Cell (PN-1000016), Chromium Single Cell A Chip Kit (PN-1000009), Chromium i7 Multiplex Kit (PN-120262). Sequencing subsequently revealed that only approximately 0.3 - 0.4 ×10^4^ cells were recovered after loading, probably due to inaccuracies in cell counting.

### Deep sequencing

#### Single-cell sequencing

Single-cell antibody repertoire libraries were sequenced on an Illumina NovaSeq 6000 system (Illumina RTA Version: V3.4.4) using a 26 x 91 bp read configuration.

#### Mammalian display library sequencing

Samples of V_H_ libraries for deep sequencing were prepared as described previously *(38)*. Briefly, genomic DNA was extracted from 1×10^6^ cells using a PureLink™ Genomic DNA mini Kit (Thermo, Cat: K182001). Extracted DNA was PCR amplified with a forward primer binding to the end of the Strep-Tag linker and a reverse primer annealing to the intronic region directly 3’ of the J segment. PCRs were performed with Q5® High-fidelity DNA polymerase (NEB, M0491L) plus GC buffer with the following reaction conditions: 1 cycle of 98°C for 30 sec; 20 cycles of 98°C, 60°C for 20 sec, 72°C for 30 sec; and a final extension at 72°C for 1 min; followed by 10°C for temporary storage. Products were checked on an agarose gel and cleaned-up through a PCR-clean up kit (DNA Clean and Concentrator-5, Zymo, D4013), followed by a secondary PCR to add illumina indexes (SFig. 4B). PCR products were then gel purified (2% TAE).

### Immunoinformatics, statistical analysis and plots

#### Data preprocessing

Raw sequencing results were processed with CellRanger (v3.1.1.0, 10x Genomics). Assembled and filtered contigs were subsequently re-aligned using Immcantation’s pRESTO/IgBlast pipeline (https://hub.docker.com/r/kleinstein/immcantation, v3.0.0, *(32)*. Sequences were aligned to the IMGT database. Only cells with a productive heavy and light chain were considered for the final analysis.

#### Statistical analysis and plots

Statistical analysis was performed using R 4.0.1 (R Core Team). Graphics were generated using the ggplot2 *(60)*, ComplexHeatmap *(61)*, and RColorBrewer *(62)* R package.

#### Enrichment

Enrichment (En) of a sequence (s) was calculated by taking the log2 of the ratio between the clonal frequency (f) in the post selection sample (sel) and the frequency observed in the background (Ab+) sample (bkg). Only sequences observed in all three (Background, S1 and S2 selected) samples are reported.

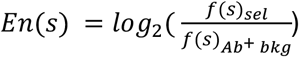

#### Overlap calculation

Overlap was determined as previously described *(63, 64)*. Briefly the number of shared sequences was divided by the size of the smaller repertoire

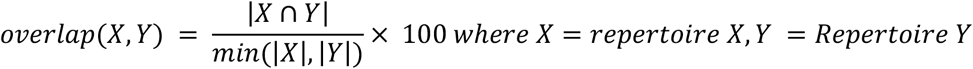

#### Sequence similarity networks

Network plots were generated using the igraph package [v.1.2.4.2, *(65)*] as previously described *(66)*, using the layout option ‘layout-nicely’ with edges drawn between amino acid sequences closer than edit distance Edit distances were calculated using the R package stringdist [v.0.9.6.3,*(67)*] using the method “hamming”.

### Design of Homology-directed Repair (HDR) Donors

Single-ORF antibody sequences were designed as reported previously *(39, 57)*. Briefly, selected paired VJ-IgK/IgL and VDJ were linked through 57–amino acid glycine-serine linker containing three tandem Strep-Tag II motifs. A universal leader sequence was added as well as 250-350 bp homology arms to the 5’ and 3’ end.

### Mammalian Cell Culture and Transfections

PnP hybridoma cells were cultured and maintained as previously described *(27, 28)*. An iteration of the established Cas9^+^ PnP-mRuby cell line was generated by flanking and deleting the mRuby-polyA gene with a pair of guides (crRNA1 and 2), as well as providing a bridging ssODN at 500 pmols to facilitate the integration of a new guide sequencing as well as an XhoI restriction site for detection of HDR on single-cell clones (**SFig. 4A**).

ssODN Sequence (*homology arms*, <u>insert</u>, **XhoI restriction site**): *GAGCCTATGGTAGTAAATACAGGCATGCCCACACTGTGAAAACAACATATGACTCCTTC*<u>GA**CTCGAG**GGT</u>*CAGGTCTTTATTTTTAACCTTTGTTATGGAGTTTTCTGAGCATTGCAGACTAATCTTGGA*

crRNA1: TCTGGGACTGTGGAGAAGAC

crRNA2: TTAAAAATAAAGACCTGGAG

crRNA3: ATATGACTCCTTCGACTCGA

Electroporation was performed with the 4D-Nucleofector™ System (Lonza) using SF Cell line 4D-Nucleofector^®^ X Kit L or X Kit S (Lonza, V4XC-2024, V4XC-2032) with the program CQ-104. Previous to nucleofection, cells were centrifuged at 150xg for 5 minutes and washed once with pre-warmed (37 °C) Opti-MEM® I Reduced Serum Medium (Thermo, 31984-062) at 1 mL per 1×10^6^ cells. The resulting cell pellet was then resuspended in SF buffer according to the manufacturer’s description, after which Alt-R gRNA and (3.5-4.5 pmoles per 1×10^6^ cells) of HDR donors were added. Cells were analyzed for HDR events approx. 3-4 days post transfection.

### Flow cytometry

Transfected cells were sorted for HDR+ by staining for Strep-Tactin-APC 1:100 (IBA lifesciences, Cat: 6-5010-001) and anti-hIgG AF488 1:100 (Jackson ImmunoResearch, Cat: 109-545-003). Enriched cells were then selected for antigen binding by incubating with SARS-CoV-2 S1-mFc 1:75 (Sinobiological, Cat: 40591-V05H1), S2-mFc 1:75 (Sinobiological, Cat: 40590-V05B) on ice for 20 min, followed by a single wash and a secondary staining with anti-mFc IgG1: PE and/or APC [clone RMG1-1] at 1:100 (Biolegend, Cat: 406608/406610). After enrichment for S1, cells were also tested for binding to SARS-CoV-2 RBD-mFc 1:375 (Sinobiological, Cat: 40592-V05H), and the same secondary staining. FACS was analyzed using FlowJo X software. At least 20’000 live events were acquired per sample to ensure sufficient statistical sampling.

### Cross-specificity ELISA

Cell supernatant containing secreted antibodies was analyzed for the presence of coronavirus specific responses. First, supernatant was collected and filtered with 0.2 μm filters (Sartorius, 16534-K). 96-well microtiter plates were coated with recombinant coronavirus antigens purchased from SinoBiological: SARS2-Spike S1+S2 (40589-V08B1), SARS1-S1 (40150-V08B1), MERS-CoV Spike (40069-V08B), HCoV-229E Spike (40605-V08B), HCoV-OC43 Spike (40607-V08B), HCoV-HKU1 Spike (40606-V08B), HCoV-NL63 Spike (40604-V08B) at 8 μg/mL in PBS at 4 °C overnight. After blocking with 2% milk, 0.05% tween PBS (PBSTM) for 1 hour at RT, cell supernatants were diluted 1:2 in 2% milk PBS (PBSM) and added to the microtiter plate for 1 hour at RT. Antigen-specific responses were detected using Strep-Tactin®-HRP (iba lifesciences, #2-1502-001) diluted 1:10’000 in PBSM, followed by repeated washing and using TMB (3,3’,5,5’-tetramethylbenzidine) (1-Step™ Ultra TMB, #34028) as a substrate. The absorbance of each sample was measured at 450 nm as well as 630 nm. All samples reported here were interrogated in the same plate.

### Octet Biolayer Interferometry

#### Kinetics

Biolayer interferometry assays were performed on the Octet Red96 (ForteBio) at 25°C, shaking at 1000 rpm. Kinetics assays were performed with Streptavidin (SA) Biosensors (ForteBio, Cat-No. 18-5019) with the following steps: (0) Hydration of SA Biosensors in 1X kinetics buffer for 30 mins (ForteBio, Cat-No. 18-1105). (1) Baseline equilibration in conditioned medium diluted 1:1 with 1X kinetics buffer for 60s (2) Regeneration of sensors (3x) in 10 mM Glycine (3) Baseline for 300s (4) Loading of antigen (SARS-CoV-2 S1 Avi-tag, Cat-No. S1N-C82E8) at 100 nM for 300 s (5) Quenching/Blocking of sensors in 50 μg/mL polyclonal human IgG in 1X KB for 30s (6) Antibody binding: sensors immersed into purified antibody at 1.9 – 500 nM in 1X KB for 350 s (7) Dissociation in 1X KB for 700 s (8) Regeneration of sensors. Curve fitting was performed using the ForteBio Octet HTX data analysis software using a 1:1 model, and a baseline correction using a reference sensor.

#### Epitope Binning

Kinetic analysis of antibody pairs was performed using Streptavidin (SA) Biosensors (ForteBio, Cat-No. 18-5019) using the “binning assay” definition with the following steps: (0) Hydration of SA Biosensors in 1X kinetics buffer for 30 mins (ForteBio, Cat-No. 18-1105). (1) Baseline equilibration in conditioned medium diluted 1:1 with 1X kinetics buffer for 60s. (2) Regeneration of sensors (3x) in 10 mM Glycine (3) Baseline for 300s. (4) Loading of antigen (SARS-CoV-2 S1 Avi-tag, Cat-No. S1N-C82E8) at 41.6 nM for 45 s. (5) Quenching/Blocking of sensors in 50 μg/mL hIgG in 1X KB for 30s. (6) Association of Ab1, ACE2 (SinoBiological, Cat-No. 10108-H08H) or Buffer (1X KB) at 166 nM for 800 s, followed by (7) short dissociation step for 10 s and then (8) the association of Ab2 at 78 nM for 400 s. (9) Regeneration of sensors. Curve fitting was performed using the ForteBio Octet HTX data analysis software using a 1:1 model, and a baseline correction with a reference sensor.

### Viral Neutralization Assays

Neutralization assays were performed as described previously *(53)*. Briefly, SARS-CoV-2 spike-pseudotyped lentiviruses incorporating a transfer plasmid encoding firefly luciferase were produced from HEK293T cells. Pseudotyped virions standardized to an input producing ~100,000 RLUs were incubated with serial dilutions of recombinant antibodies for 60 min at 37 °C prior to the addition of ~15,000 HEK293T-ACE2 cells and incubation for 48h. Luminescence was measured using Bright-Glo (Promega) on a GM-2000 luminometer.

## Acknowledgements

We acknowledge the ETH Zurich D-BSSE Single Cell Unit and the Genomics Facility Basel for excellent support and assistance.

## Author Contributions

R.A.E., F.B., R.V-L. and S.T.R. designed experiments; D.M.M., B.W., F.B., R.V-L., R.B.D.R., A.P.C., F.R., M.S., assisted in patient sample collection, analysis and cell isolation experiments; R.A.E., K.L.H and C.W. performed genome editing and antibody screening experiments; D.J.S. and B.M. designed and performed viral neutralization experiments; R.E., C.R.W. and S.F. analyzed sequencing data; R.A.E., C.R.W. and S.T.R. wrote the paper, with feedback and input from all authors.

## Data and Materials Availability

The raw FASTQ files from deep sequencing that support the findings of this study will be deposited (following peer-review and publication) in the Sequence Read Archive (SRA) with the primary accession code(s) <code(s) (https://www.ncbi.nlm.nih.gov/sra)>. Additional data that support the findings of this study are available from the corresponding author upon reasonable request.

## Supplementary Materials

Figures S1 - S7

Tables S1 - S6

