## Supplementary Material for "Single-cell sequencing of plasma cells from COVID-19 patients reveals highly expanded clonal lineages produce specific and neutralizing antibodies to SARS-CoV-2"

**Supplementary Materials:**

Figures S1 - S7

Tables S1 - S6

### Supplementary Materials:

#### Supplementary Figures:

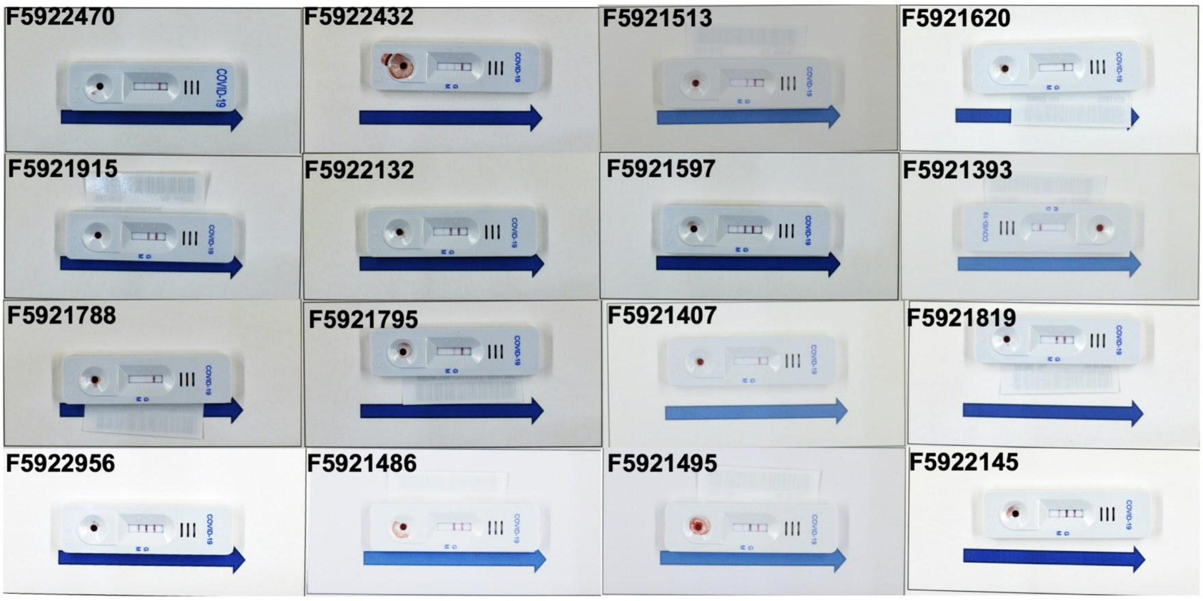


**Fig. S1. Antibody response measurements from COVID-19 patients.** Standardized images of the Hightop lateral flow assay (IgM, IgG and Control) for each patient.

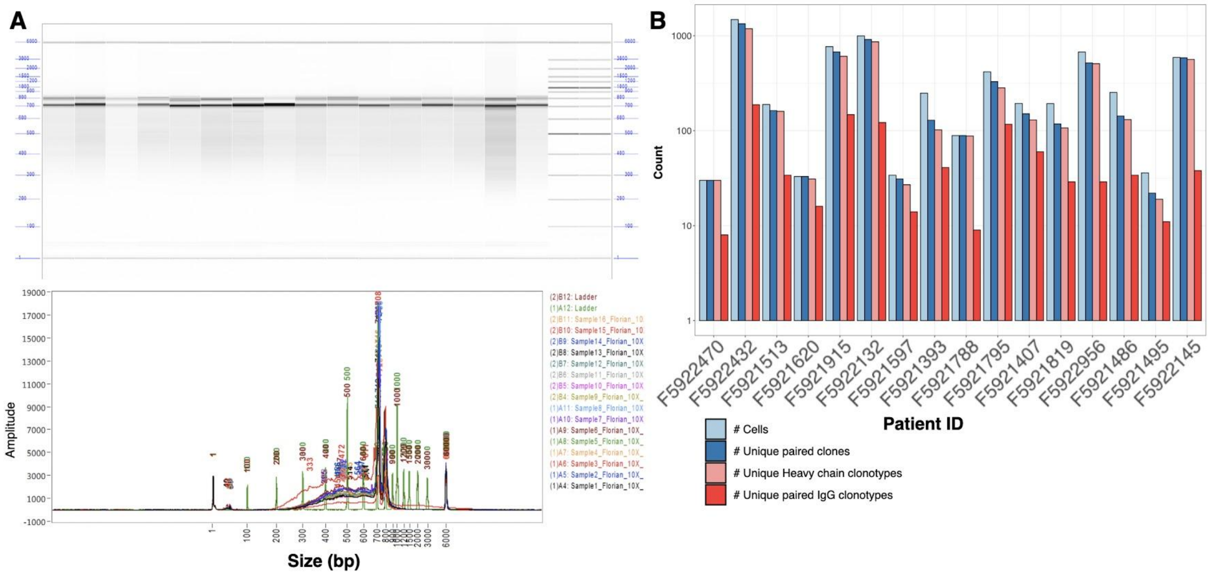


**Fig. S2A. Sequencing data quality.** (A) Library preparation QC data for 10X. (B) Sequencing statistics, read counts, cells, clonal lineages, etc.

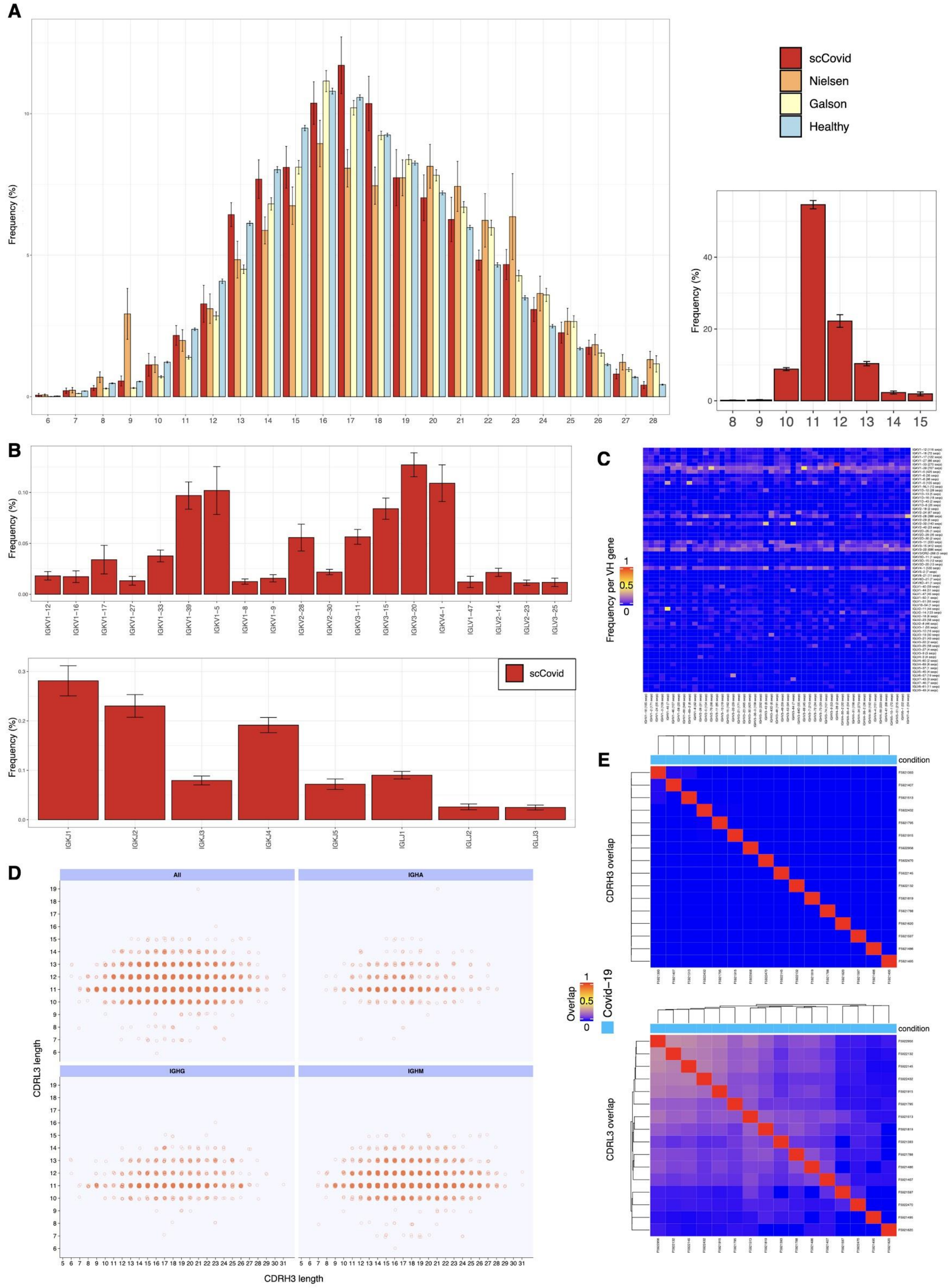


**Fig. S2B. Additional repertoire features.** (A) Length distribution of heavy and light chain CDR3 sequences. (B) Light chain germline gene usage, V (top) and J (bottom). (C) Heavy-Light chain pairing frequencies pooled across all scCovid repertoires. (D) Lengths of paired CDRH3 and CDRL3s per isotype. (E) Overlap of CDRH3 and CDRL3 sequences across patients.

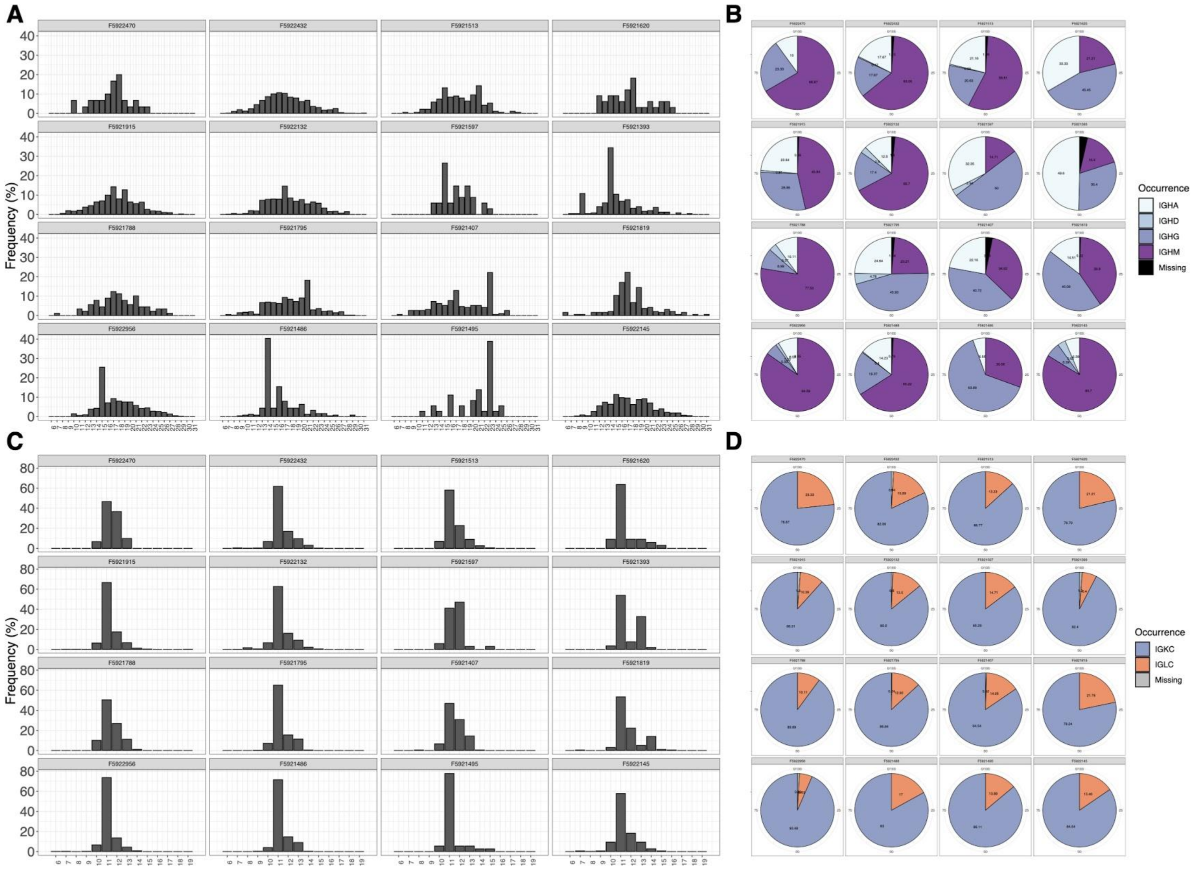


**Fig. S2C. Per sample statistics.** (A,C) Length distribution of heavy and light chain CDR3s per patient sample. (B,D) Isotype usage per patient for heavy and light chains.

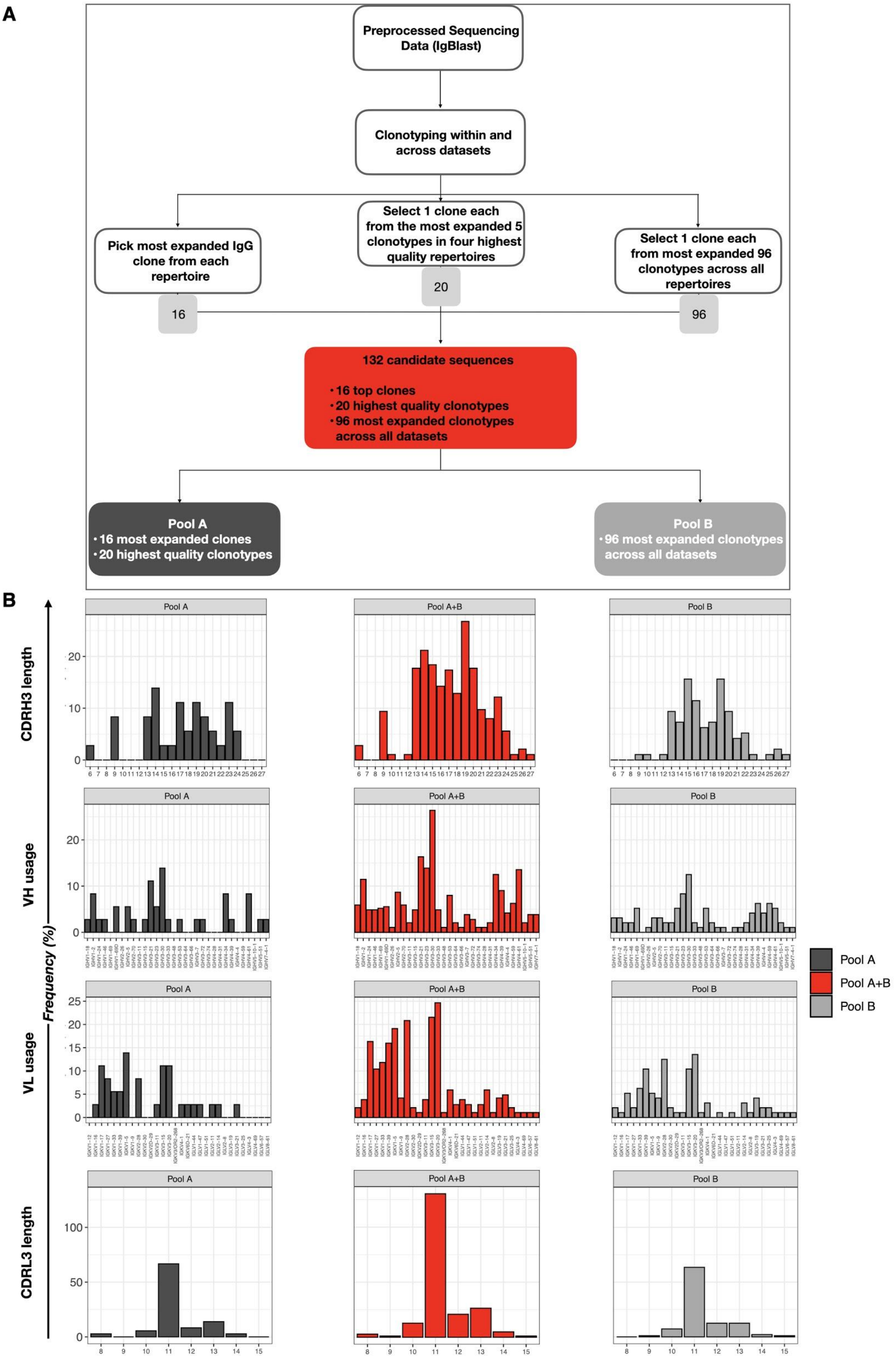


**Fig. S3. Workflow for selection of candidate sequences.** A. Candidate sequences were selected in three steps. Initially the top IgG clone in each sample was identified and selected. This was followed by the selection of 20 sequences through determining the most expanded clone within the top 5 clonal lineages observed in four high quality patient repertoires (as determined by QC and number of cells detected). Finally, one clone from the top 96 clonal lineages observed across all repertoires (not previously selected) was added to the set of selected sequences. (B) CDRH3 and -L3 length distribution for the selected clonal lineages, as well as the $V_H/V_L$ gene usage, split into the respective pools, Pool A (black), Pool B (grey) and Pool A+B (red).

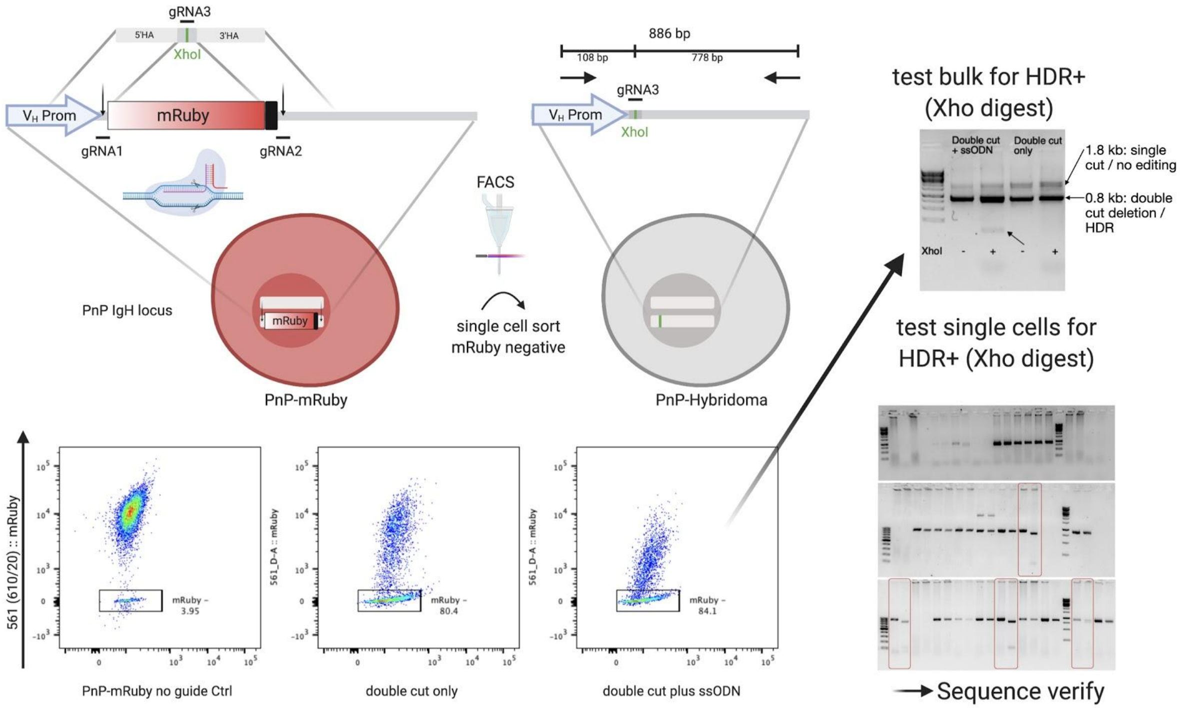


**Fig. S4A. Iteration of PnP-mRuby platform.** Given the occasional residual mRuby signal post Ab+ enrichment decreasing the efficiency of selection on other 561 nm fluorochromes such as PE, mRuby was deleted from the PnP-IgH locus. This also facilitated the integration of short homology arm constructs, increasing the HDR efficiency (Data not shown).

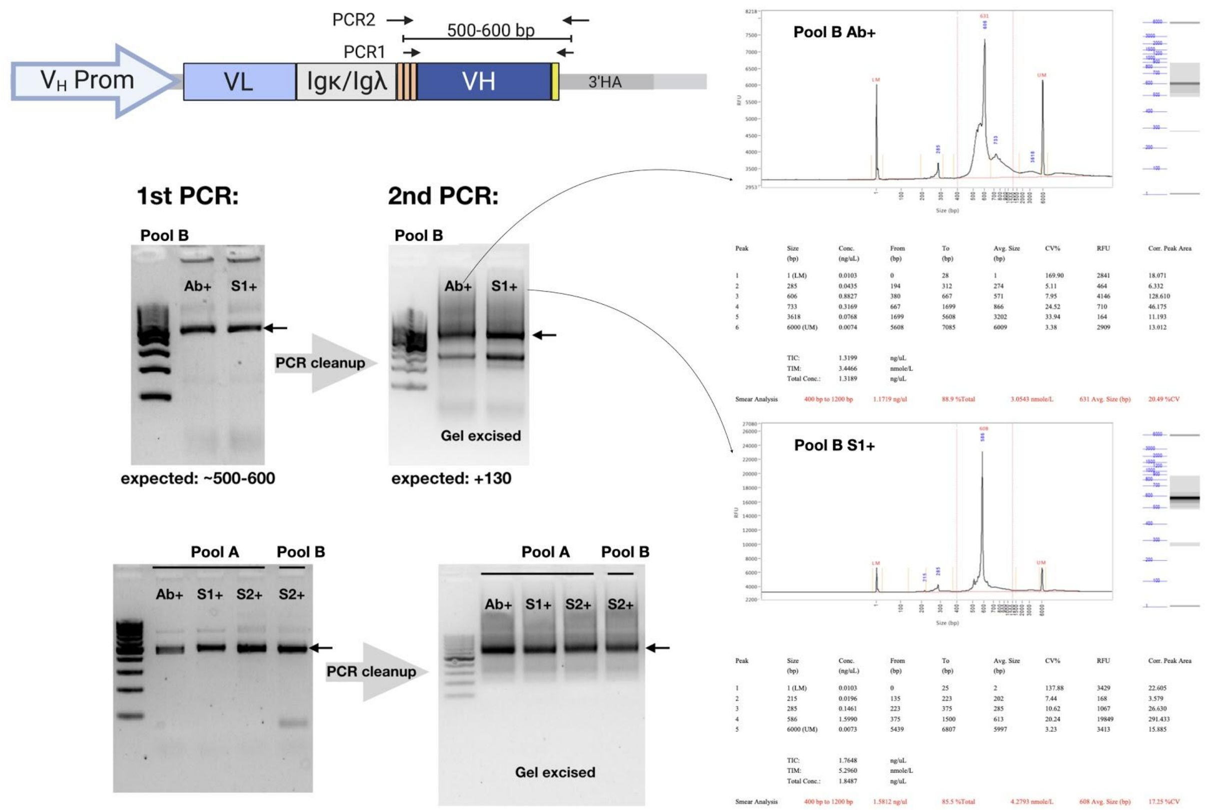


**Fig. S4B. Template amplification for deep sequencing.** The respective heavy chains of bulk antigen-binding and/or antibody displaying hybridoma were amplified in a first PCR (PCR1), adding illumina adapters to the sequence. After PCR cleanup, the fragments were reamplified in a 2nd PCR (PCR2) to add illumina indices for each respective sample, followed by a gel excision. Samples were loaded onto a fragment analyzer post excision to determine their purity. Examples are shown for Pool B Ab+ and S1+ sorted samples.

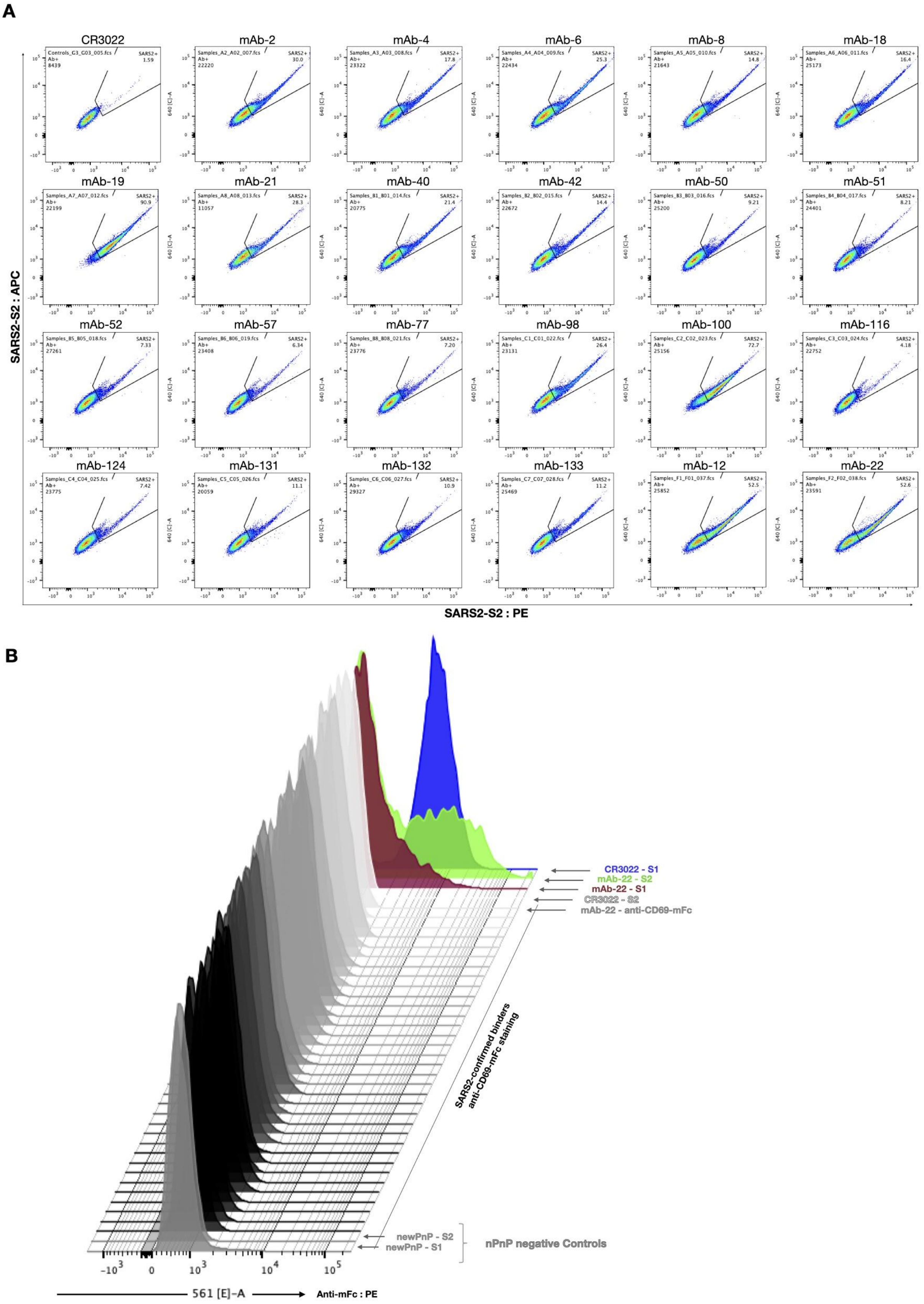


**Fig. S5. Background controls.** (A) Double staining of selected S2 binders. Low S2 binding mAbs were double stained for both SARS-CoV-2: S2-mFc plus anti-mouse IgG1:APC and :PE to check for PE related background and false positivity. (B) Background determination for unrelated antigen-mFc**.** Select SARS-CoV-2 binders were additionally stained with anti-hCD69 (mouse IgG1), followed by the previously used anti-mouse Fc secondary staining (here shown: anti-mFc : PE).

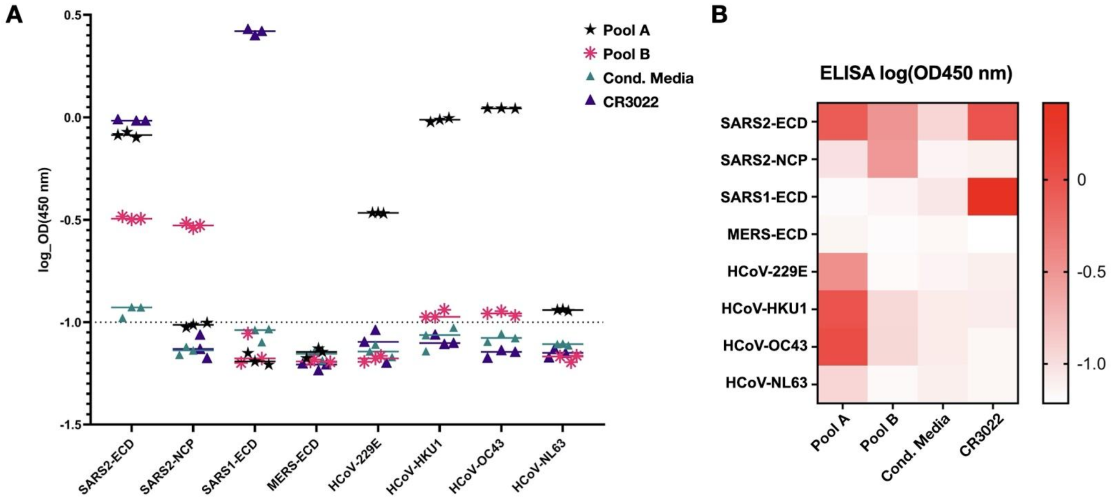


**Fig. S6. Cross-specificity ELISA to other human coronavirus antigens.** (A) Log OD450-570 nm values given for all antigens in response to Pool A or B supernatant, conditioned medium or CR3022 supernatant. Coating antigens on the X-axis. (B) Heatmap displaying the results from A. Conditioned media of the founder nPnP cell line was used to control for supernatant dependent background signal.

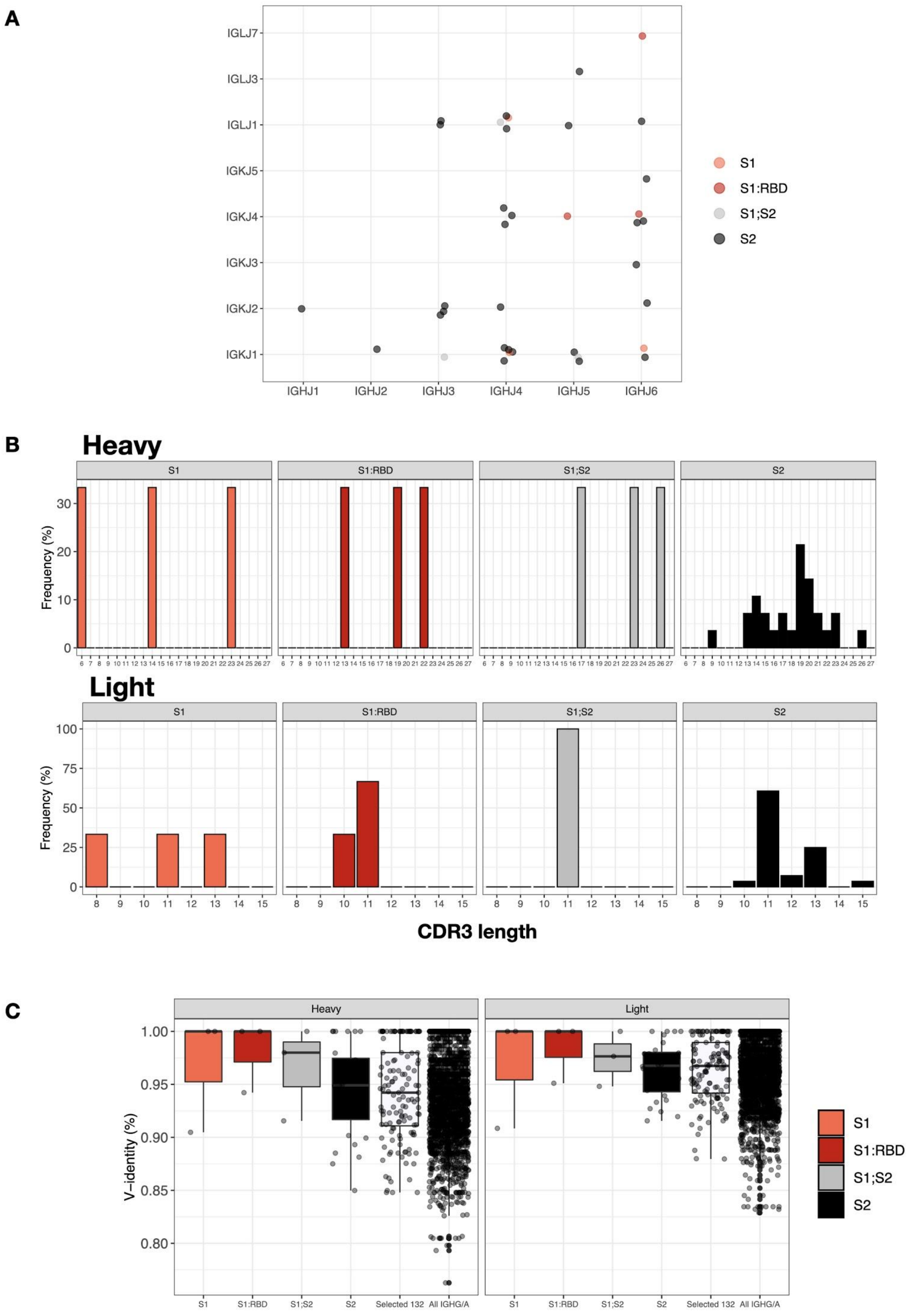


**Fig. S7. Extended sequence features of SARS-CoV-2 reactive antibodies.** (A) Length distribution of heavy and light chain CDR3s, followed by (B) V- and J-gene usage, and (C) the V-gene identity (level of somatic hypermutation) grouped according to antigen reactivity.

**Supplementary Tables:**

https://docs.google.com/spreadsheets/d/1w0V4yKs2OAuPJiXi0iwpkIML2LruAvZaQ1UeLlwdI9I/edit?ts=5fd110b8#gid=2117901290

**Table S1.** CellRanger metrics. Resulting sequencing reads and cell numbers determined from CellRanger v3.1.1.0, for all patient samples.

**Table S2.** 132 candidate sequences were selected. Fully annotated sequences of the selected sequences.

**Table S3.** Enrichment of selected variants. The 132 selected variants were examined for enrichment.

**Table S4.** Subset of confirmed SARS-CoV-2 Binders. The top 50% of binders include 7 S1 binders (3 of which also bind RBD), 9 S2 binders and one S1;S2 cross-reactive antibody.

**Table S5.** Relating reactive sequences back to patients. Three categories of patients: No reactive sequences identified, Only S2 reactive and both S1,S2 reactive (37/132 selected, ~28% were reactive) (mean hits/selected per repertoire 35%).

**Table S6.** Sequence similar hits in databases. Binders in comparison with sequences found in the CovAbDab.
